# Bora Bridges Aurora-A Activation and Substrate Recognition of PLK1

**DOI:** 10.1101/2025.08.08.668882

**Authors:** Jennifer A Miles, Matthew Batchelor, Martin Walko, Vanda Gunning, Andrew J Wilson, Megan H Wright, Richard Bayliss

## Abstract

Cell cycle transitions are orchestrated by regulated protein kinase activities. The activation of PLK1 in late G2 is a decisive step for cells to enter mitosis, requiring phosphorylation of its activation loop by Aurora-A, facilitated by the intrinsically disordered protein Bora. Despite its biological importance, the structural basis of this mechanism has remained unresolved. Here, we present models of the Aurora-A/Bora complex and the ternary Aurora-A/Bora/PLK1 complex, validated with site-specific mutagenesis, biochemical assays and NMR spectroscopy. Bora wraps around the N-lobe of Aurora-A, occupying the pockets used by its other activators. A CDK1 phosphorylation site on Bora (Ser112) mimics the structural role of Aurora-A activation loop phosphorylation within a TPX2-like binding motif. In the ternary complex, Bora bridges the two kinases, orienting the activation loop of PLK1 towards the active site of Aurora-A. Bora residues 56-66 form a critical interface with a conserved pocket on the PLK1 C-helix that is analogous to the TPX2-binding Y-pocket of Aurora-A. Aurora-A phosphorylation of Bora Ser59 creates an additional interaction that increases the efficiency of PLK1 phosphorylation. These findings deepen our understanding of how Aurora-A activity is fine-tuned by its disordered binding partners and establish a mechanistic framework for its Bora-dependent activation of PLK1.

## Introduction

Protein kinases such as Aurora-A, PLK1 and CDK1 have critical roles in orchestrating mitosis, especially in the regulation of the G2/M transition and mitotic spindle assembly and function. Aurora-A is a member of the Aurora family of kinases, that also includes kinetochore-associated Aurora-B and Aurora-C. It has roles in mitotic spindle assembly, DNA repair, centrosome maturation and cilia regulation (Willems *et al*, 2018). Structurally, Aurora-A contains a disordered N-terminal domain (aa 1-121), followed by a kinase domain (aa 122-387) and a short disordered C-terminal region (aa 388-403). Canonical activation of its kinase function requires phosphorylation of Thr288 in the activation loop, a process that primarily occurs by autophosphorylation. Phosphorylated Aurora-A is enriched on spindle poles and is also found along spindle microtubules close to the poles (Ohashi *et al*, 2006; Holder *et al*, 2024). Aurora-A activity is tightly regulated by activators and inhibitors, in particular through the binding of intrinsically disordered proteins that ‘complete’ the incomplete Aurora-A core kinase domain (Levinson, 2018; Bayliss *et al*, 2012). Activator proteins, such as TPX2 (Bayliss *et al*, 2003), can stimulate Aurora-A autophosphorylation on Thr288, leading to an active kinase.

PLK1 is a member of the Polo-like kinase family first identified in *Drosophila melanogaster* which is comprised of 5 members in humans (Llamazares *et al*, 1991; Sunkel & Glover, 1988; Korns *et al*, 2022). PLK1 is composed of an N-terminal kinase domain linked to two tandem polo-box domains (PBD). The PBDs are required for recognition of protein substrates and control of PLK1 localisation (Park *et al*, 2010). PLK1 is frequently overexpressed in cancer and is linked to a poor prognosis (Eckerdt *et al*, 2005). Activation of PLK1 is achieved through the phosphorylation of Thr210 by Aurora-A kinase (Macůrek *et al*, 2008; Seki *et al*, 2008a; Chan *et al*, 2008) during the G2/M transition following successful DNA damage repair (Seki *et al*, 2008b). Active PLK1 then phosphorylates and activates key mitotic regulators such as the phosphatase CDC25 and the ubiquitin ligase APC/C (Moshe *et al*, 2004; Qian *et al*, 2001).

Bora is an intrinsically disordered protein of 559 amino acids that was discovered in *Drosophila melanogaster* (Hutterer *et al*, 2006) and its overexpression has been observed in human bladder and colorectal cancer, and adenocarcinoma samples (Cheng *et al*, 2020; Mahajan *et al*, 2023; Zhang *et al*, 2017). The selective modification of PLK1 by Aurora-A on Thr210 is mediated by Bora (Seki *et al*, 2008b; Macůrek *et al*, 2008; Hutterer *et al*, 2006). Bora has a high content of serine (15%) and threonine (6.6%) residues (Thomas *et al*, 2016) and a total of fourteen sites are phosphorylated by CDK1–Cyclin A *in vitro* and twelve *in vivo* (Feine *et al*, 2014; Thomas *et al*, 2016). Three of the sites that are modified by CDK1–Cyclin A (Ser41, Ser112, Ser137) are evolutionarily conserved and important for the function of Bora, including in *C. elegans* as well as human cells (Thomas *et al*, 2016; Tavernier *et al*, 2015; Parrilla *et al*, 2016; Pintard & Archambault, 2018).

Phosphorylation of Bora promotes its interaction with PLK1, which is diminished when three N-terminal phosphorylation sites are mutated (S41A, S112A, S137A) (Thomas *et al*, 2016). The roles of other phosphorylations have yet to be defined, although Thr52 phosphorylation may be required for degradation of Bora (Feine *et al*, 2014).

The precise mechanism by which PLK1, Aurora-A and Bora come together to bring about Thr210 phosphorylation is unclear. However, it most likely involves a transient ternary complex, as observed by cross-linking mass spectrometry (Lössl *et al*, 2016). The mechanism does not require canonical activation of Aurora-A on Thr288, but does require Bora that is phosphorylated on Ser112, which can act in *trans* to mimic activation loop phosphorylation (Figure 1A) (Tavernier *et al*, 2021). It is not known how the Aurora-A/Bora complex interacts with PLK1 to catalyse its phosphorylation on Thr210.

**Figure 1.**
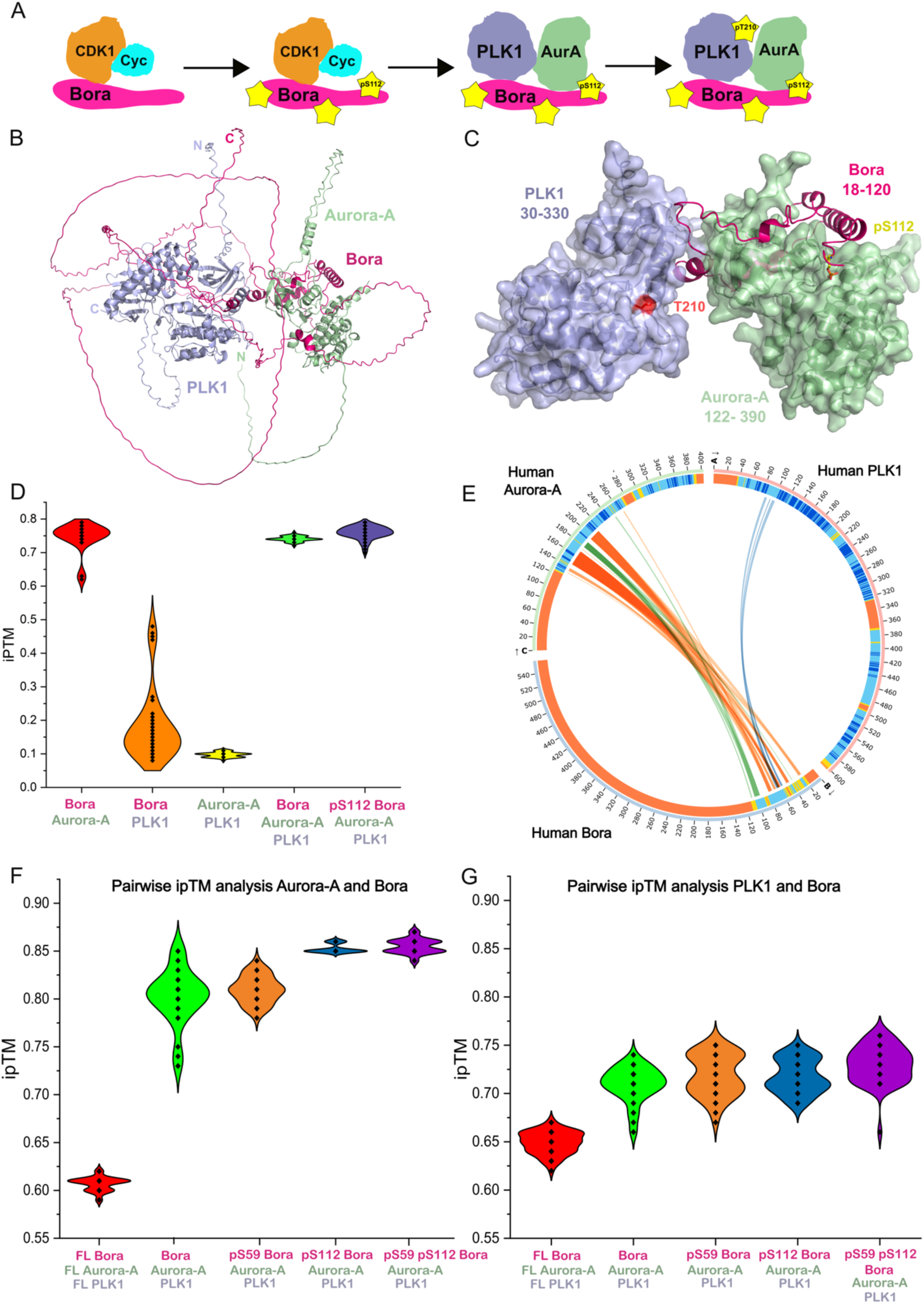
Bora Bridges Aurora-A Activation and Substrate Recognition of PLK1. **A:** Model showing the role of Bora in the facilitation of PLK1 phosphorylation by Aurora-A kinase. Bora is phosphorylated on numerous sites, including Ser112 by CDK1–Cyclin A. Phosphorylated Bora then binds more tightly to Aurora-A and acts in *trans* to activate the unphosphorylated Aurora-A, resulting in the phosphorylation of PLK1 at Thr210 (human numbering). **B:** AlphaFold3 model of the ternary complex between full-length human Aurora-A, PLK1 and Bora. Aurora-A 1-403 is shown in green, with PLK1 1-603 in blue and Bora 1-559 in magenta. **C:** AlphaFold3 model of the truncated ternary complex between human Aurora-A, PLK1 and Bora. Aurora-A 122-390 is shown in green, with PLK1 30-330 in blue and Bora 18-120 in magenta. Threonine 210 in PLK1 that is modified by Aurora-A is highlighted in red, with the phosphorylated Serine 112 in human Bora highlighted in yellow. **D:** Violin plot of iPTM scores from 50 models of the truncated binary and ternary complexes (Bora 18-120, Aurora-A 122-403 and PLK1 30-330) produced by AlphaFold3 (using alphafoldserver.com). **E:** AlphaBridge representation of the ternary complex between human Aurora-A, PLK1 and Bora. The sequences are coloured based on the confidence in the modelling of these regions, with blue indicating high confidence and orange indicating low confidence. The residues predicted to be interacting between the three proteins are linked with curved lines. **F:** Violin plot of iPTM scores between Bora and Aurora-A in the ternary complexes of Bora, Aurora-A and PLK1 produced by AlphaFold3 (using alphafoldserver.com). The phosphorylation state of Bora is listed. FL = full-length, with all the other complexes being modelled with the truncated proteins (Bora 18-120, Aurora-A 122-403 and PLK1 30-330). **G:** Violin plot of iPTM scores between Bora and PLK1 in the ternary complexes of Bora, Aurora-A and PLK1 produced by AlphaFold3 (using alphafoldserver.com). The phosphorylation state of Bora is listed. FL = full-length, with all the other complexes being modelled with the truncated proteins (Bora 18-120, Aurora-A 122-403 and PLK1 30-330).

High-resolution, experimental structural studies on transient interactions such as those involving PLK1 and Aurora-A remain challenging. We therefore took advantage of recent advancements in computational modelling that enable accurate predictions of the structures of proteins and their complexes (Bryant *et al*, 2022; Baek *et al*, 2021), and specifically AlphaFold3 which has the capability to include phosphorylated side-chains, ligands and ions (Abramson *et al*, 2024). We report a high-confidence structural model for the complex between PLK1, Aurora-A and Bora. The model was validated using structure-guided mutagenesis, biochemical assays and NMR spectroscopy. It provides a rationale for the roles of Bora phosphorylation in the interaction, including an additional site modified by Aurora-A that we characterised.

## Results

### Bora is a bridge between Aurora-A and PLK1 in the ternary complex

The complex of the three full-length human proteins (Bora (1-559), Aurora-A (1-403), PLK1 (1-603)) was modelled using AlphaFold2 (Mirdita *et al*, 2022) and AlphaFold3 (Abramson *et al*, 2024) (Figure 1B, AlphaFold3 model with Bora shown in magenta, Aurora-A in green and PLK1 in blue). The model had an overall interface predicted template modelling (ipTM) score of 0.63 using both AlphaFold3 and AlphaFold2 (Supplementary Table 3). The ipTM score is a measure of the accuracy of the predicted relative positions of the residues in a complex, on a scale of 0-1. Bora was modelled as highly disordered, with residues 20-113 wrapped around the Aurora-A kinase domain. Residues 52-73 of Bora are modelled between the PLK1 kinase domain and Aurora-A kinase domain, with residues 58 to 68 predicted to form an alpha-helix. Unsurprisingly for an intrinsically disordered protein, the confidence of most of the Bora structural prediction was low, particularly over residues 175-559 (Supplementary Figure 1A).

The model was simplified by removal of the low confidence regions of all three proteins, whilst preserving the key interactions. Bora 18-120 was wrapped around Aurora-A, forming the ternary interface with PLK1, and a short region of Bora (245-257) that includes the phosphorylated Ser252 site interacted with the PBD of PLK1 (Supplementary Figure 1B) (Chan *et al*, 2008). The PBD is located on the opposite side of the PLK1 kinase domain to the Bora interface. We therefore focussed on the kinase domains of both PLK1 (30-330) and Aurora-A (122-390), and the minimal region of Bora (18-120) required to stimulate Aurora-A (Tavernier *et al*, 2021). Models based on these truncated sequences perfectly conserved the interactions observed in the full-length protein model with Bora forming the core of the ternary interface (Figure 1C, Bora in magenta between Aurora-A in green and PLK1 in blue). The average ipTM generated from 50 models produced with AlphaFold3 was 0.74 (Figure 1D, plotted in green). Analysis of the ternary complex with AlphaBridge (Álvarez-Salmoral *et al*, 2024) identified two interfaces, between Aurora-A (130-280) and Bora (21-110), and Bora (56-66) with PLK1 (86-103) (Figure 1E). There is no direct interface between Aurora-A and PLK1, and so Bora can be considered as a bridge between the two kinases. Models generated using both AlphaFold2 and AlphaFold3 of the ternary complex were consistent (Supplementary Figure 2A and B).

Modelling the binary complexes produced consistent, high confidence models of Aurora-A bound to Bora but more variable, lower confidence models of PLK1 bound to either Bora or Aurora-A (Figure 1D, Table S3). A similar trend was observed in the pairwise ipTM scores for the interfaces in the ternary complexes (Figure 1F,G). All models of PLK1/Aurora-A complexes were of very low confidence.

Bora wraps around the N-lobe of Aurora-A in all the models of the Aurora-A/Bora binary complex (AlphaBridge interface summary in Supplementary Figure 1C and superposed models in Supplementary Figure 2C). In contrast, without Bora being ‘tethered’ in place by Aurora-A, the models of a complex between the PLK1 kinase domain and Bora 18-120 are highly variable (Supplementary Figure 3A and B). When Bora is limited to just residues (52-72) that are predicted to bind to PLK1, these models are more consistent, placing the Bora sequence at the same site in 9 out of 10 models (Supplementary Figure 3C, Phe56 and Trp58 shown in magenta).

The interface on PLK1 predicted to interact with Bora is analogous to the Y-pocket on Aurora-A, which interacts with Tyr8 and Tyr10 in TPX2 (Bayliss *et al*, 2003; McIntyre *et al*, 2017a). Since this pocket in PLK1 is predicted to interact with a phenylalanine and tyrosine in Bora, this site will be provisionally labelled as the ‘FW pocket’. Previous mass spectrometry analysis identified peptides within this region as contributing to the interaction between human PLK1 and human Bora (Bora peptides 50-78, 58-78 (Seki *et al*, 2008a)).

### Biochemical validation of the ternary complex model

Our model suggests that the interaction between Bora and PLK1 is mediated by a small section of Bora that interacts with the ‘FW’ pocket in PLK1 (Figure 2A). A fluorescently labelled version of this short region of Bora (52-73) showed weak binding to the PLK1 kinase domain in a fluorescence polarisation assay (Figure 2B, black, a K82R PLK1 mutant was used to increase the stability of the protein). The interaction was substantially reduced in a variant of PLK1 with changes to the ‘FW’ pocket (R106A, S99R) (Figure 2B, red).

**Figure 2.**
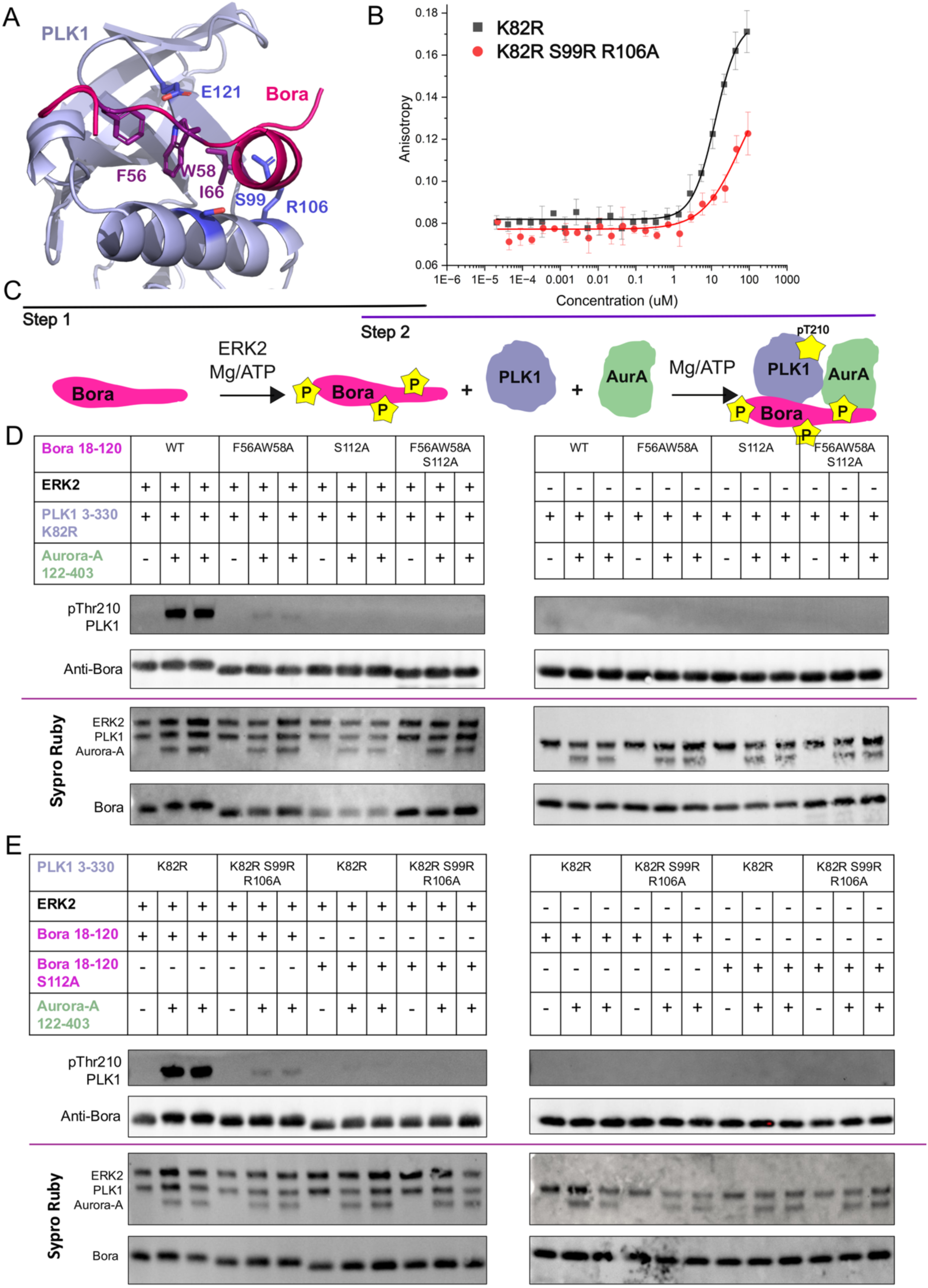
Biochemical validation of the Bora:PLK1 interface. **A:** Cartoon representation of the predicted interface between PLK1 kinase domain (blue) and Bora 18-120 (magenta). **B:** Fluorescence anisotropy based direct binding assay looking at the interaction of FAM Bora 53-72 with PLK1 3-330 (K82R kinase dead version, shown in black) and PLK1 mutated in the predicted interaction interface (K82R S99R R106A shown in red). **C:** Schematic of the PLK1 phosphorylation assay, where Bora 18-120 is preincubated with ERK2 and ATP to produce Bora phosphorylated before Aurora-A and PLK1 are added in the second step. **D:** Western blot analysis of the levels of phosphorylation of PLK1 kinase domain K82R at Thr210 when different Bora mutants are preincubated with ERK2 and added to wild-type dephosphorylated Aurora-A kinase domain (122-403). Inputs are shown with Sypro Ruby staining of nitrocellulose membranes. **E:** Western blot analysis of the levels of phosphorylation of PLK1 kinase domain K82R and the S99R R106A mutant at Thr210 when Bora wild-type or Bora S112A is preincubated with ERK2 and added to wild-type dephosphorylated Aurora-A kinase domain. Inputs are shown with Sypro Ruby staining of nitrocellulose membranes.

The model was validated further using an *in vitro* assay with phosphorylation of Thr210 of PLK1 as a readout. Mutations were introduced into Bora 18-120 to remove two of the hydrophobic residues that are predicted to interact with PLK1 (Phe56 and Trp58, Figure 2A). Purified mutated Bora was then pre-phosphorylated with ERK2, before the addition of wild-type unphosphorylated Aurora-A kinase domain and PLK1 3-330 K82R as the substrate (Figure 2C). ERK2 kinase was chosen as it is selective for (S/T)P motifs equivalent to those phosphorylated *in vivo* by CDK1 but can phosphorylate this shorter sequence of Bora which lacks the Cy-motif needed to recruit the CDK1–Cyclin complex (Tavernier *et al*, 2021). When WT Bora was pre-phosphorylated by ERK2, phosphorylation of PLK1 at Thr210 was stimulated by Aurora-A (Figure 2D, lanes 2 and 3) (Tavernier *et al*, 2021). There was a significant reduction in PLK1 phosphorylation at Thr210 when the pre-phosphorylated F56A/W58A mutant of Bora was used (Figure 2D, lanes 5 and 6). A similar effect was seen when Ser112 in Bora was mutated to alanine (Figure 2D, lanes 8 and 9) and when the mutations were combined (Figure 2D, lanes 11 and 12). Unexpectedly, the F56A/W58A Bora was less efficiently phosphorylated on S112A (Supplementary Figure S11, F compared to H and Supplementary Table S4). To mitigate for this, the assay used a 5-fold molar excess of Bora compared to levels of PLK1 and Aurora-A to ensure that enough phosphorylated Bora was present. A Bora variant in which only S112 can be phosphorylated by ERK2 (Bora S27A S41A T52A, labelled as ‘ERK’ mutant) also lead to clear stimulation of Aurora-A activity, which was also abrogated upon inclusion of the F56A/W58A mutation (Supplementary Figure 4A).

There was a significant reduction in levels of phosphorylation on PLK1 at Thr210 when the ‘FW’ pocket mutant of PLK1 (S99R R106A) was used as a substrate (Figure 2E, comparing lanes 2 and 3 with lanes 5 and 6). These mutations did not impact the overall structure of PLK, as the ‘FW’ variant retained interaction with a DARPin that binds to the kinase domain (Supplementary Figure 4B and C) (Bandeiras *et al*, 2008).

We conclude that the interaction of the region of Bora centred on Phe56/Trp58 is critical for its interaction with PLK1, and for the subsequent phosphorylation of Thr210 by Aurora-A.

### Conservation of Bora-PLK1 interaction site

Examination of the predicted interface between PLK1 and Bora revealed a pattern of conserved residues (Figure 3A-D). Orthologues of the human sequences were identified using a PSI-Blast search and the sequences aligned using MAFFT (Supplementary Figures 5 and 6). The residues that are predicted to interact with Bora in PLK1 were selected using a PDBe PISA (Krissinel & Henrick, 2007) analysis of the model of the ternary complex. Lys86, Leu89 and Arg95 at the interface are well conserved in PLK1 (human numbering, Figure 3B). Furthermore, the two hydrophobic residues that point into the FW pocket (Phe56 and Trp58) are conserved between all Bora orthologues (Figure 3D), as is the proline at the end of the short helix in Bora in the complex (Pro68). This suggests that the interaction between Bora and PLK1 using this interface is likely to be similar in many organisms. To assess this hypothesis, AlphaFold3 was used to model the orthologues from an organism distant in evolution from humans, *Strongylocentrous purparatus* (sea urchin). The truncated ternary complex has a very similar arrangement to the human ternary complex (Figure 3E compared to Figure 1C). The interface between PLK1 and Bora is conserved, with Phe55 and Trp57 from Bora pointing into the surface of PLK1 (Figure 3F).

**Figure 3.**
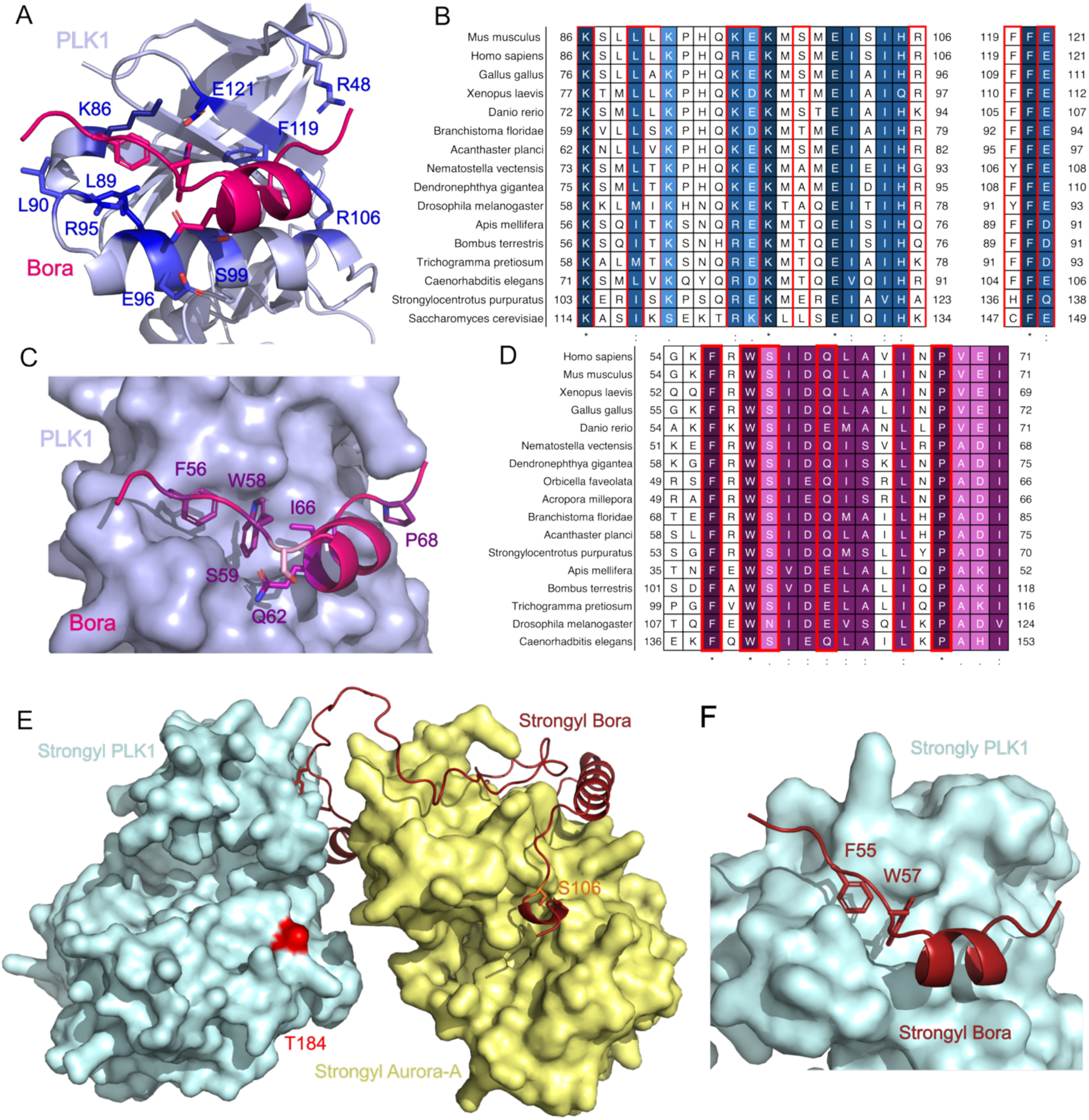
Conservation of the Bora:PLK1 interface. **A:** Model of Bora (magenta) interacting with the FW pocket on PLK1 (blue), with residues in the interface from the PLK1 side coloured based on conservation. The darker the residue, the more conserved it is. **B:** Selected regions of the MAFFT sequence alignment of the PLK1 kinase domain from diverse species. The residues highlighted in red are at the predicted interface with Bora. **C:** Model of Bora (magenta) interacting with the FW pocket on PLK1 (blue), with residues in the interface from the Bora side coloured based on conservation. The darker the residue, the more conserved it is. **D:** Sequence alignment of the region of Bora that is predicted to interact with PLK1. The darker the residue, the more conserved it is. The residues highlighted in red are at the predicted interface with PLK1. **E:** AlphaFold3 model of the ternary complex from *Strongylocentrotus purpuratus* with PLK1 shown in cyan, Bora shown in dark red and Aurora-A in yellow. Threonine 184 that is predicted to be phosphorylated by Aurora-A is shown in bright red on the surface of PLK1, with the phosphorylated serine in Bora shown in orange. **F:** Highlighting the conserved interface between PLK1 (cyan) and Bora (red) in *Strongylocentrotus purpuratus* orthologues, showing Phe55 and Trp57 in the pocket on the surface of PLK1.

### Characterisation of Bora 1-120 and its interaction with Aurora-A using NMR

^15^N-^13^C labelled Bora 1-120 was expressed and purified prior to characterisation by NMR. The ^1^H-^15^N HSQC spectrum shows features of a disordered protein as seen previously (Tavernier *et al*, 2021), albeit there is enough peak dispersion to suggest the sequence may contain some elements of order (Figure 4A, spectra in black). The ^1^H-^15^N HSQC was straightforwardly assigned using triple resonance experiments (Supplementary Figure 7). Two regions in the C-terminal half of Bora 1-120 (Pro73-Arg87 and Lys90-Thr105) have significant helical propensity (regions exhibit positive C⍺ and CO secondary shifts and negative Cβ secondary shifts, Supplementary Figure 8A-D). Regions with weaker helical propensities are also observed in the N-terminal half (Tyr31-Thr38 and Ile60–Val65). The helical propensity regions in the unbound Bora NMR data match up very well with the helical regions of Bora present in the Aurora-A/Bora and Aurora-A/Bora/PLK1 AlphaFold models. This finding supports the AlphaFold models and indicates that the interaction builds upon latent structure within the ‘disordered’ Bora chain.

**Figure 4.**
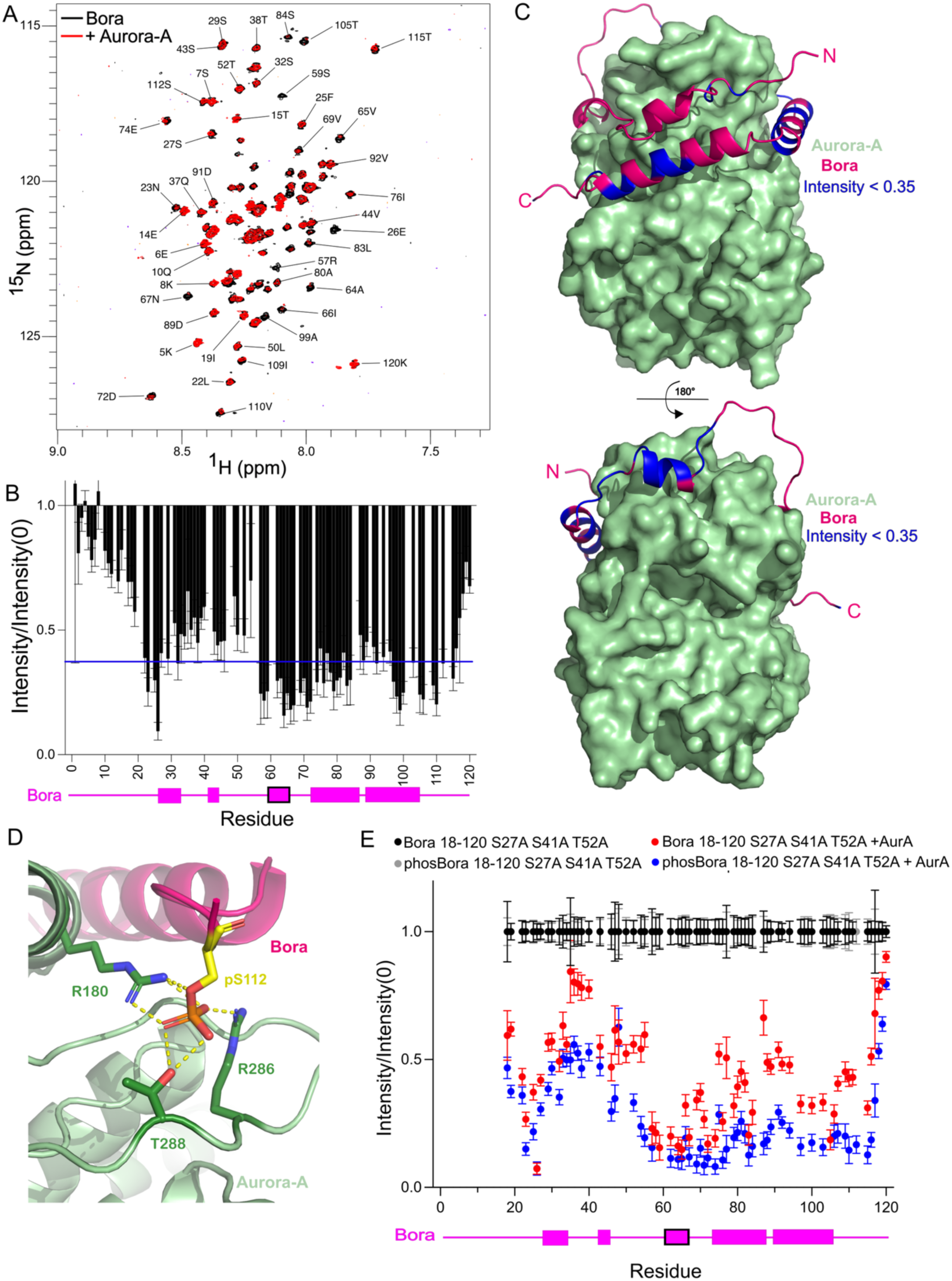
NMR analysis of Bora interaction with Aurora-A. **A:** ^1^H-^15^N HSQC recorded on *Homo sapiens* Bora 1-120 in the absence (black) and presence of human Aurora-A 122-403 (red). A significant number of Bora peaks disappeared in the presence of Aurora-A. **B:** The intensity changes of residues in Bora 1-120 in the absence and presence of Aurora-A 122-403 ([Bora]:[AurA] = 2:1). The model-predicted secondary structure of Bora in the complex is represented below in pink, with predicted alpha helices shown as rectangles. The helix predicted to interact with the FW pocket of PLK1 is bordered in black. **C:** Mapping the residues involved in binding onto the predicted structure of Bora 1-120 bound to Aurora-A. Aurora-A is shown in green, with Bora represented in magenta. Residues in Bora that show a significant loss in peak volume upon inclusion of Aurora-A are shown in blue (intensity change below 0.35). **D:** Analysis of the AlphaFold3 model of Bora phosphorylated at Ser112 (yellow) bound to Aurora-A (shown in green). Phosphorylated Ser112 in Bora is predicted to co-ordinate two arginine residues in Aurora-A as well as Thr288 in the activation loop of Aurora-A. **E**: Change in peak intensity in the ^1^H-^15^N HSQC spectrum of ^15^N-labelled Bora 18-120 S27A S51A T52A with and without phosphorylation of Bora by ERK at Ser112 (black - no phosphorylation, grey - with phosphorylation by ERK) and when Aurora-A is present (red - unphosphorylated Bora and Aurora-A, blue - phosphorylated Bora with Aurora-A). The reduction in peak intensities around the Ser112 (and at other parts of the sequence) is higher for the phosphorylated version. The model-predicted secondary structure of Bora in the complex is represented below in pink, with predicted alpha helices shown as rectangles. The helix predicted to interact with the FW pocket of PLK1 is bordered in black.

Residues in the two high-helical-propensity regions stand out from the rest of the Bora sequence by having elevated ^15^N *R*_2_ relaxation rates (and thus, comparatively low peak intensities at 10 °C, Supplementary Figure 8E and G) and elevated hetNOE ^15^N-^1^H values (Supplementary Figure 8F). These two regions also maintain or increase their ^1^H-^15^N HSQC peak intensities on increasing temperature, whereas intensities for residues elsewhere within Bora 1-120 tend to reduce with temperature (Supplementary Figure 8H). The region of Bora predicted to interact with both Aurora-A and PLK1 (close to Ile60) has similarly elevated relaxation features. The hetNOE features in the C-terminus indicate deviation away from the more freely dynamic, disordered behaviour seen in the N-terminal half of Bora 1-120, to areas with restricted motion on the ps–ns timescale from partial helix formation. The elevated *R*_2_ values could also contain a contribution from some slower ms–ms dynamics in these regions.

When Aurora-A 122-403 (kinase domain) was titrated into ^15^N-^13^C labelled Bora 1-120, no clear chemical shift perturbations were observed, but loss in peak intensities to the point of disappearances were seen at positions all through the sequence (Figure 4A, shown in red). Peak intensity changes seen across the Bora sequence at a [Bora]:[AurA] molar ratio of 2:1 are shown in Figure 4B. The Bora residues with the most significant loss in peak intensity, which are likely to be those most constrained by the interaction, when mapped onto the AlphaFold3 model of Aurora-A/Bora complex (Figure 4C, blue) match very well with those predicted to most closely interact. By contrast, and as observed previously (Tavernier *et al*, 2021), titrating PLK1 into labelled Bora resulted in very little by way of spectral change even at higher molar ratios, highlighting the transient nature of the Bora/PLK1 interaction.

### Phosphorylation at Ser112 stabilises the interaction between Aurora-A and Bora

AlphaFold3 supports the modelling of post-translational modifications (PTMs) such as phosphorylation. When the truncated ternary complex between Bora, Aurora-A and PLK1 was modelled with phosphorylation of Bora at Ser112, a site which has been shown to act in *trans* to activate Aurora-A (Tavernier *et al*, 2021), the ipTM score was slightly improved (Figure 1D, shown in dark blue, average ipTM 0.76 compared to 0.74). The phosphoryl group at this site is predicted to interact with Arg180, Arg286 and Thr288 in Aurora-A (Figure 4D). This is comparable to how phosphorylated Thr288 in the activation loop of Aurora-A is part of a stabilised activation loop when TPX2 is bound (Bayliss *et al*, 2003).

The effect of pSer112 in the pairwise ipTM scores were also analysed using 10 models produced with AlphaFold3 (5 models from each of 10 runs). There is on average a 0.05 uplift in ipTM score when looking at the model of Aurora-A bound to pSer112 (Figure 1F, shown in blue). Whereas the presence of the phosphorylation of on Bora pS112 doesn’t affect the pairwise Bora-PLK1 ipTM score significantly (Figure 1G, shown in blue). This is consistent with a specific effect of pSer112 on stabilisation of the interaction with Aurora-A.

To further investigate the role of this phosphorylation *in vitro*, we phosphorylated ^15^N labelled Bora 18-120 S27A S41A T52A (ERK mutant) with the kinase ERK. In ^1^H-^15^N HSQC spectra, ERK phosphorylation of Bora 18-120 S27A S41A T52A specifically targets Ser112 (Supplementary Figure S9A). When Aurora-A kinase domain is titrated into this phosphorylated Bora, we see a similar profile but with larger intensity losses in the pSer112 region of Bora, suggesting a stronger local interaction with Aurora-A (Figure 4E). As well as a decrease in intensity around pSer112 in Bora, see we an overall effect with decreased intensity across most of the Bora sequence. Potentially the increased binding at pSer112 acts as a tether holding Aurora-A and Bora together, improving the stability of the interaction overall. This model is in agreement with the observed higher affinity of Aurora-A for phosphorylated Bora (Tavernier *et al*, 2021).

Crystal structures are available of Aurora-A in complex with several other binders: TPX2, CEP192, TACC3 and N-Myc (Bayliss *et al*, 2003; Richards *et al*, 2016; Holder *et al*, 2024; Burgess *et al*, 2018; Park *et al*, 2025). In the AlphaFold models Bora is predicted to exploit the same set of pockets on the N-lobe of Aurora-A as TPX2, CEP192 and TACC3 using similar interactions (Figure 5A). In the overlaid structures, Bora Phe25 overlaps with Phe19 in TPX2 and Phe490 in CEP192 (Figure 5B), and Bora Phe103/Phe104 overlaps with Trp34 and Phe35 from TPX2 (Figure 5E). Phe45 in Bora predicted to interact with the Y-pocket in Aurora-A, although it does not closely resemble Tyr8 and Tyr10 of TPX2 at this site (Figure 5C). Ile71 of Bora is predicted to bind in the F-pocket on the other side of the Aurora-A N-lobe that can be occupied by either Phe525 of TACC3 or Phe508 of CEP192 (Figure 5D).

**Figure 5.**
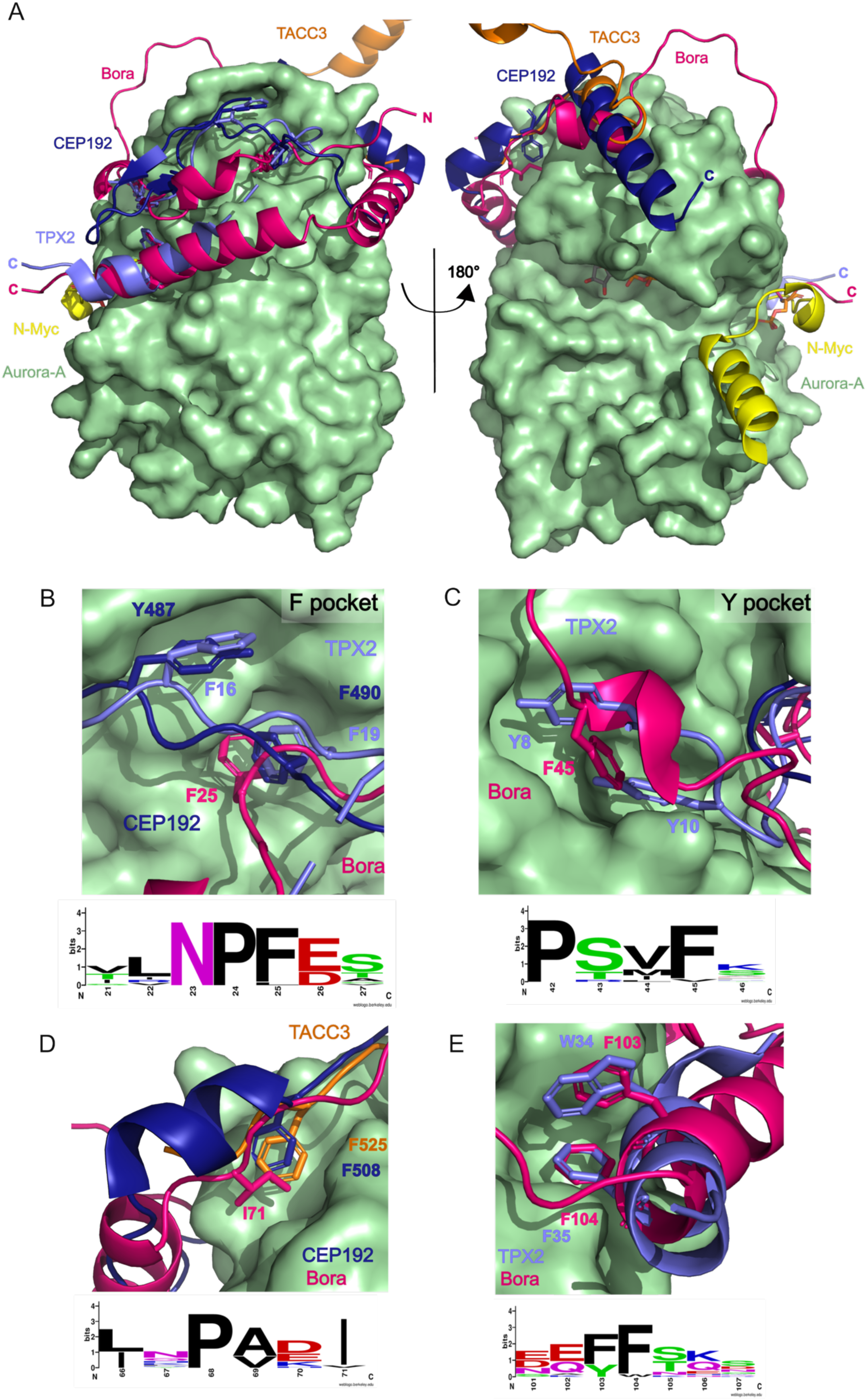
Structural comparison of Aurora-A complexes. **A:** Comparison of the model of Bora (magenta) bound to Aurora-A (green) with the crystal structures of Aurora-A complexes. TPX2 1-43 is shown in light blue (PDB 1OL5), CEP192 468-533 in dark blue (PDB 8PR7), N-Myc 61-89 in yellow (PDB 5G1X) and TACC3 in orange (PDB 5ODT). **B:** Comparisons of predicted interaction of Bora with Aurora-A, focusing on the F-pocket of Aurora-A. F25 of Bora is predicted to bind in the F pocket on the surface on Aurora-A, which can also be occupied by F19 in TPX2 or F490 in CEP192. The conservation of these sites in Bora is shown as a WebLogo below. **C:** Comparison of the predicted interaction of Bora with Aurora-A in the Aurora-A ‘Y-pocket’. Bora F45 is predicted to bind to Aurora-A at this site, utilising the pocket that is occupied by Y8 and Y10 in TPX2. The conservation of these sites in Bora is shown as a WebLogo below. F45 in Bora is highly conserved. **D:** Comparison of the predicted interaction of Bora with Aurora-A, focusing on the pocket at the top of Aurora-A that both TACC3 and CEP192 interact with. TACC3 and CEP192 have phenylalanine residues that interact with a pocket on the top of the N-lobe of Aurora-A (F525 and F508, respectively). The modelling predicts that I71 from Bora occupies this site on Aurora-A. The conservation of these sites in Bora is shown as a WebLogo below. **E:** Comparison of the predicted interaction of Bora with Aurora-A, focusing on the region near the activation loop. F103 and F104 of Bora are predicted to interact with Aurora-A similarly to how TPX2 W34 and F35 interact with the surface of Aurora-A. The conservation of these sites in Bora is shown as a WebLogo below.

An NMR-based competition assay was used to probe the Aurora-A/Bora interaction sites in solution. ^1^H-^15^N HSQC spectra of ^15^N-^13^C labelled Bora 1-120 were again recorded before and after the addition of Aurora-A, resulting in a decrease in the intensities across the Bora sequence (Figure 6A). Next, Aurora-A binding proteins with known binding sites were added (TPX2 1-43, CEP192 442-533) and spectra re-measured, resulting in the partial rescue of intensities (Figure 6B, 6C). This indicates that TPX2 and CEP192 can compete for Aurora-A-binding with Bora – liberating the Bora leads to peak intensity recovery. The recovery on addition of TPX2 was most marked in the region around Bora Phe25, and CEP192 was more effective for peak recovery of Bora’s C-terminal region. These observations are consistent with the predicted binding sites of Bora compared to those utilised by TACC3, CEP192 and TPX2 on Aurora-A (Figure 6D and E).

**Figure 6.**
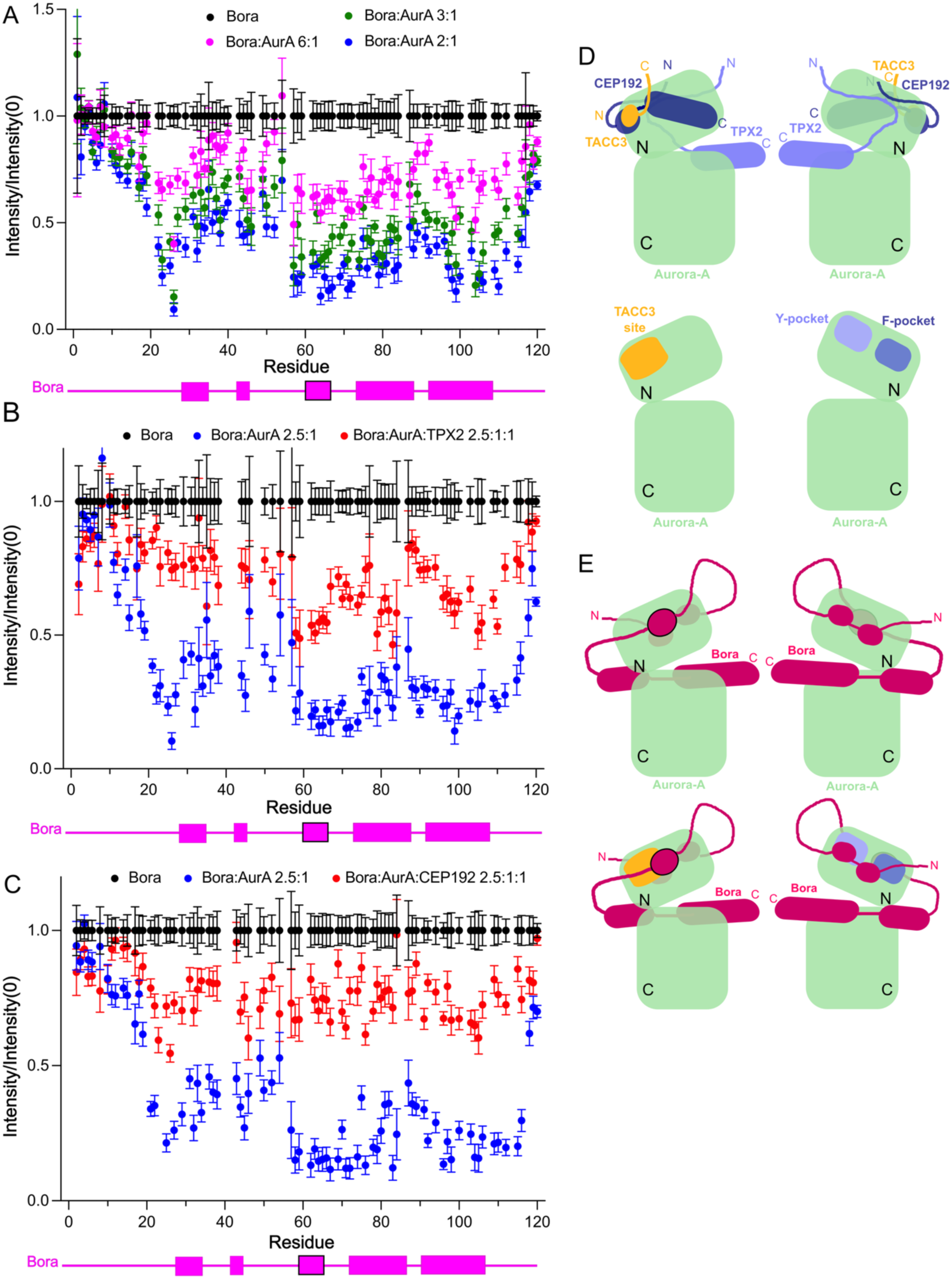
NMR-based competition assays for binding sites on Aurora-A. **A:** The relative change in intensity for peaks assigned to residues in Bora 1-120 in the presence of increasing amounts of Aurora-A kinase domain. Interaction with a binding partner leads to changes in the chemical environment and relaxation/dynamic behaviour, resulting in changes in peak position and intensity, respectively; the largest changes are typically observed at the binding site(s). In this case, Aurora-A interaction led to peak intensity loss at sites of interaction but no clear chemical shift perturbations. The model-predicted secondary structure of Bora in the complex is represented below in pink, with predicted alpha helices shown as rectangles. The helix predicted to interact with the FW pocket of PLK1 is bordered in black. **B:** Aurora-A competition assay of Bora 1-120 with TPX2 1-43. Labelled Bora 1-120 is incubated with Aurora-A (2.5:1 molar ratio), leading to position-dependent peak intensity loss as the Bora:Aurora-A interactions occur (as per (**A**) shown in blue). Introduction of unlabelled TPX2 1-43 (1:1 molar ratio with Aurora-A) then leads to rescue of the Bora signal as TPX2-binding reduces Bora:Aurora-A interactions. Recovery of peak intensity is most effective close to the N-terminal binding site close to Phe25. **C:** Aurora-A competition assay of Bora 1-120 with CEP192 468-533. Labelled Bora 1-120 is incubated with Aurora-A (2.5:1 molar ratio), leading to position-dependent peak intensity loss as Bora:Aurora-A interactions occur (as per (**A**) shown in blue). Introduction of unlabelled CEP192 (1:1 molar ratio with Aurora-A) then leads to rescue of the Bora signal as CEP192-binding reduces Bora:Aurora-A interactions. Here, recovery of peak intensity is more effective for binding sites in the C-terminal half of Bora 1-120. **D:** Illustration of the sites used by TACC3 (orange), TPX2 (light blue) and CEP192 (dark blue) to interact with Aurora-A (light green). The TACC3 binding site, Y pocket and F pocket are highlighted on the Aurora-A N-lobe. **E:** Illustration of the predicted sites used by Bora (magenta) to bind to Aurora-A (light green). The TACC3 binding site, Y pocket and F pocket are highlighted on the Aurora-A N-lobe. The helix predicted to interact with the PLK1 ‘FW’ pocket is bordered in black in the images on the left.

### Aurora-A phosphorylation of Bora Ser59 enhances PLK1 activation

Given that Bora is heavily phosphorylated by numerous kinases, we considered whether modification of any site other than Ser112 could influence the interaction with PLK1 or Aurora-A. A peptide array covering the Bora sequence was used to identify sites that can be phosphorylated by Aurora-A (Figure 7). Bora peptides containing Ser59 exhibited the most significant increase in staining (Figure 7A). This site has previously been identified, with 100% modification observed after 20 min of Aurora-A incubation with Bora (Lössl *et al*, 2016). This residue is conserved in most species, but not *Drosophila melanogaster* (Figure 4D), where an asparagine is present at this position. The consensus sequence for substrate phosphorylation by Aurora-A is R/K/N-R-X-S/T-B, with B denoting any hydrophobic residue with the exception of Pro (Ferrari *et al*, 2005). This is similar to the sequence in Bora around Ser59 (F-R-W-**S**-I).

**Figure 7.**
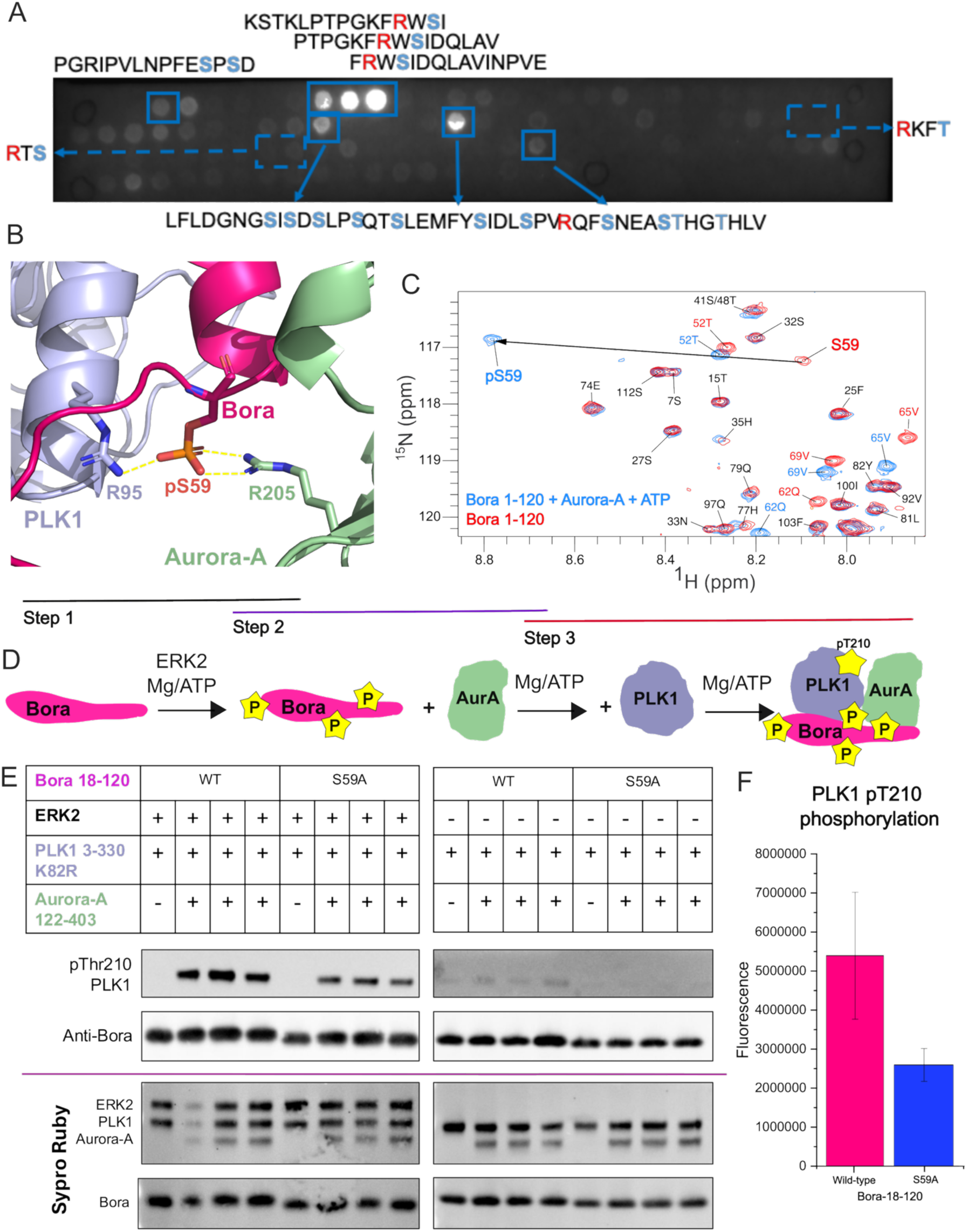
Aurora-A phosphorylated Bora at Ser59. **A:** Peptide array of Bora divided into peptides of 15 amino acids. The array was incubated with ATP and active Aurora-A kinase domain before probing with Pro-Q Diamond Phosphoprotein stain and imaging. Clear phosphorylation is seen on the peptides that include Ser59 of Bora. **B:** Modelling of pSer59 in the ternary complex of Bora (magenta), Aurora-A (green) and PLK1 (blue). The phosphorylated residue is in the interface between PLK1 and Aurora-A. Interactions are predicted between the phosphate on Ser59 and Arg95 in PLK1 and Arg205 in Aurora-A. **C:** ^1^H-^15^N HSQC spectrum of ^15^N-labelled human Bora 1-120 in the absence (red) and presence (blue) of active Aurora-A kinase domain and ATP. A clear shift in the position of the Ser59 peak to the left indicates that this is it being modified by Aurora-A *in vitro*. **D:** Altered assay schematic where Aurora-A wild-type kinase domain is preincubated with phosphorylated Bora 18-120 wild-type and S59A before the PLK1 K82R kinase domain is included as a substrate. **E:** Western blot analysis of the levels of phosphorylation of PLK1 kinase domain at Thr210 when Bora 18-120 wild-type and S59A are preincubated with ERK2, followed by Aurora-A before addition to PLK1 K82R kinase domain. Blots were probed with antibodies to Bora and pThr210 in PLK1. The samples were repeated in triplicate to allow quantification. Inputs are shown with Sypro Ruby staining of nitrocellulose membranes. **F:** Quantification of the levels of phosphorylation of PLK1 at Thr210 in the presence of either wild-type Bora 18-120 or S59A Bora.

The predicted effect of Bora phosphorylation at Ser59 to the interaction with Aurora-A and PLK1 in the predicted ternary complex was modelled using AlphaFold3. The model indicates that phosphorylation of Ser59 makes an additional interaction with Arg95 of PLK1 and Arg205 of Aurora-A (Figure 7B). This may stabilise a ternary complex further to facilitate PLK1 phosphorylation.

The phosphorylation of ^15^N-labelled human Bora 1-120 was monitored using NMR. Clean, specific phosphorylation at Ser59 was observed through a substantial downfield ^1^H shift in the peak for this residue when Bora 1-120 was incubated with Aurora-A (Figure 7C, full spectra in Supplementary Figure 9B). Ser59 phosphorylation resulted in chemical shift perturbations (CSPs) for residues Thr52-Ile71, and an unusual upfield ^15^N shift for Ser59. This may indicate an increase in local helical propensity in this region of Bora, consistent with the helical conformation predicted in the ternary complex model.

To probe the functional relevance of Ser59, it was mutated to alanine and the variant protein used in an *in vitro* assay of PLK1 phosphorylation at T210 (Figure 7D). Following incubation with phosphorylated Bora S59A with wild-type Aurora-A 122-403, PLK1 phosphorylation after 30 min showed a 50% reduction compared to using wild-type Bora (Figure 7E and F). This indicates that Bora Ser59 phosphorylation mediated by Aurora-A increases the efficiency of PLK1 phosphorylation.

## Discussion

It has proven challenging to determine experimental structures of kinase-substrate complexes, and only thirty have been deposited in the PDB, eleven of which are kinase autophosphorylation structures (Faezov & Dunbrack, 2023). The mechanism by which Aurora-A recognises PLK1 via Bora has also eluded experimental structure determination, due to disfavourable properties of the system: Bora is a disordered protein that requires phosphorylation on specific sites to interact with and activate Aurora-A, and the PLK1/Bora interaction is transient, like most kinase-substrate pairs (Bruinsma *et al*, 2015; Tavernier *et al*, 2021).

Here we have used AlphaFold3 to model how Bora, Aurora-A and PLK1 come together to phosphorylate PLK1 on Thr210. Bora is predicted to form a bridge to bring the Aurora-A and PLK1 together, through interactions with pockets on the surfaces of the N-lobes of the kinases. A critical interaction is formed between a motif in Bora (56-66) and a pocket at the C-helix of PLK1. Phosphorylation of Bora on Ser112 is important for the phosphorylation of PLK1 as it mimics the structural role of Aurora-A activation loop phosphorylation in the context of a TPX2-like binding motif and in the context of unphosphorylated Aurora-A. Aurora-A phosphorylation of Bora Ser59 also enhances the efficiency of PLK1 phosphorylation. Ser59 is a good substrate for Aurora-A, is highly conserved, and lies within the critical motif of Bora positioned at the interface with PLK1 in the ternary complex. The predicted interface between Bora and the PLK1 kinase domain is small (buried surface area of 574 ± 39 Å^2^ with unphosphorylated Bora, 598 ± 42 Å^2^ with pS59 pS112 Bora), consistent with a weak interaction. In contrast, the predicted interface between Aurora-A and Bora is large (buried surface area of 2739 ± 181 Å^2^ with unphosphorylated Bora and 2663 ± 126 Å^2^ with pS59 pS112 Bora).

One limitation of the modelling is that, although the activation loop of PLK1 faces the active site of Aurora-A, Thr210 is too distant for a productive phospho-transfer reaction. There is some flexibility in the PLK1 position, indicating that movement of PLK1 Thr210 towards the active site is possible, but it would not be sufficient to close the gap. This is because the activation loop of PLK1 adopts a closed, active-like conformation, not an open/extended conformation that would be needed to act as a substrate. Attempts to model PLK1 with a more extended activation loop conformation, using a range of templates, were unsuccessful. AlphaFold3 is ill-suited for predicting dynamic features of the complex, as it has been trained against stable, experimental structures in the PDB, in which transient conformations of kinase-substrate complexes are underrepresented.

The Aurora-A/Bora interaction was previously proposed to resemble that of the TPX2/Aurora-A interaction, based on the sequence similarities of two motifs. Motif one, comprising 22-35, was proposed to resemble TPX2 7-20, but whereas TPX2 binds the Y-pocket via Tyr8 and Tyr10, the sequence alignment of Bora with TPX2 placed only Phe25 of Bora into the Y-pocket of Aurora-A, preceded by an “NP” sequence that does not resemble the “YS” of TPX2. In the AlphaFold3 model of the ternary complex, the chain direction of Bora is opposite to that of TPX2, and Phe25 of Bora is predicted to interact with the F-pocket in Aurora-A. Motif two in the previous study, comprising Bora 100-111, was predicted to bind to Aurora-A like TPX2 31-42. This hypothesis is supported by the AlphaFold3 model, in which Phe103 and Phe104 of Bora are structurally equivalent to Trp34 and Phe35 of TPX2. Both TPX2 and Bora appear to stabilise the activation loop of Aurora-A, either by facilitating interactions between a phosphorylated residue in the activation loop (Thr288) and the rest of Aurora-A, or by acting in *trans* and providing a phosphate to support this stabilisation (pSer112 in Bora).

Crystal structures of the chromosome passenger complex protein INCENP bound to the Aurora-B and C kinase domains show that it too wraps around the N-lobe, and phosphorylation of Ser893 and S894 in the INCENP sequence is vital for the complete stimulation of kinase activity (Abdul Azeez *et al*, 2019; Elkins *et al*, 2012). The structure of INCENP bound to Aurora-C (PDB 6GR8) was aligned with the model of Aurora-A bound to Bora, highlighting three points of similarity in these interfaces (Supplementary Figure 10). Short helices in INCENP and Bora bind above the kinase C-helix via similar hydrophobic interactions, namely Phe45 of Bora compared with Phe881 in INCENP (Supplementary Figure 10B). Another similarity is the helix that is predicted to wrap around the N-lobe (Supplementary Figure 10C). Bora I76 and Q79 overlay on top of INCENP I855 and Q585 respectively, suggesting Bora has a very similar mode of binding to this region of the kinase domain of Aurora proteins. Finally, near the activation loop, the phosphorylated Ser893 and Ser894 in INCENP are positioned at a similar site to the pS112 in Bora, with S893 and S894 interacting with the activation loop of Aurora-C (Supplementary Figure 10D). Thus, the way that Bora interacts with Aurora-A combines features similar to both INCENP and TPX2.

The identification of selective kinase inhibitors remains a significant challenge (Karaman *et al*, 2008). We have shown that the FW pocket on PLK1 mediates its interaction with Bora. This pocket is structurally analogous to the Y-pocket of Aurora-A and the PDK1-interacting fragment (PIF) pocket in PDK1 – both known regulatory sites that influence kinase activation and selectivity (Biondi *et al*, 2000; Balendran *et al*, 1999). Ongoing efforts to develop selective binders for these pockets (Rettenmaier *et al*, 2014; Engel, 2013; Arencibia *et al*, 2017; Stockwell *et al*, 2024; McIntyre *et al*, 2017b) suggest that targeting the FW pocket could similarly disrupt PLK1 activation, offering a potential route to a new class of specific PLK1 inhibitors for PLK1– dependent cancers.

## Methods

### Reagents

Human His-GST tagged ERK2 (2-360) and ERK1 (2-379) were purchased from MRC PPU Reagents and Services (DU650, DU1509). Human Bora 18-120, 18-120 S27A S41A T52A and PLK1 binding DARPin were purchased as codon optimised sequences from GenScript. Oligonucleotides used for site-directed mutagenesis and subcloning were ordered from IDT and are detailed in the supplementary table 1. The site-directed mutagenesis was performed using the Q5 site-directed mutagenesis kit (NEB). The constructs used in this publication are listed in supplementary table 2. All DNA constructs were verified by sequencing. Pro-Q Diamond PhosphoProtein gel stain was purchased from ThermoFisher Scientific.

The antibodies used in this study are: anti-Bora (Santa Cruz Ca. sc-393741), antiPhospho-Plk1 pT210 (Cell Signalling Technologies Cat. No. 5472), goat anti-rabbit StarBright 700 secondary (Biorad Cat. No. 12004162), goat anti-mouse StarBright 700 secondary antibody (Biorad Cat. No. 12004159).

### Protein production

Bora 18-120, 1-120 and mutants were cloned into the petSUMO vector with an N-terminal TEV protease cleavable His-SUMO tag. This was transformed into B834 RIL cells. 4 litres of cells were inoculated with 10 ml of overnight culture. Protein expression was induced overnight with 0.5 mM IPTG at 20 °C. The pellet was resuspended in 10 ml per litre of ice-cold lysis buffer (50 mM TRIS pH 7.5, 500 mM NaCl, 10% glycerol, 0.5 mM TCEP, 20 mM imidazole, EDTA-free protease inhibitors). The cells were sonicated for 10 sec on, 20 sec off for 4 min 10 sec total at 60%. The soluble fraction was collected at 17000 rpm for 45 min. This was then filtered through 0.45 μm filters before loading onto a HisTrap FF. The bound protein was eluted in a gradient of 500 mM imidazole. This was dialysed overnight into 250 mM NaCl, 50 mM TRIS pH 7.5, 1 mM TCEP, 10% glycerol in the presence of TEV protease. The following morning the protein was incubated with 3 ml of Ni-NTA equilibrated in dialysis buffer on the roller for 45 min to bind any uncut protein or His-SUMO. The flowthrough containing Bora was concentrated and loaded onto an SD200 16/600 size exclusion column equilibrated in 300 mM NaCl, 50 mM TRIS pH 7.5, 10% glycerol, 1 mM TCEP. The clean fractions were concentrated in a 10 kDa cut off concentrator and the protein was flash frozen and stored at −80 °C.

Human PLK1 3-330 and mutants were cloned into the petSUMO vector with an N-terminal TEV protease cleavable His-SUMO tag. This was transformed into B834 RIL cells. 4 litres of cells were inoculated with 10 ml of overnight culture. Protein expression was induced overnight with 0.5 mM IPTG at 20 °C. The pellet was resuspended in 10 ml per litre of ice-cold lysis buffer (50 mM TRIS pH 7.5, 500 mM NaCl, 10% glycerol, 0.5 mM TCEP, 20 mM imidazole, 5 mM MgCl_2_, EDTA-free protease inhibitors). The cells were sonicated for 10 sec on, 20 sec off for 4 min 10 sec total at 60%. The soluble fraction was collected at 17000 rpm for 45 min. This was then filtered through 0.45 μm filters before loading onto a HisTrap HP. The bound protein was eluted in a gradient of 500 mM imidazole. This was dialysed overnight into 250 mM NaCl, 50mM TRIS pH 7.5, 1 mM TCEP, 10% glycerol, 5 mM MgCl_2_ in the presence of TEV protease to remove the His-SUMO tag. The following morning the protein was loaded onto a HisTrap HP equilibrated in dialysis buffer, and the flow through collected. Bound protein was eluted in a gradient of imidazole after washing with 4 CVs of dialysis buffer. The PLK1 partly eluted in the flow through and partly in the start of the imidazole gradient. This was concentrated in a 30 kDa cut off concentrator and loaded onto the SD200 16/600 size exclusion column equilibrated in 300 mM NaCl, 50 mM TRIS pH 7.5, 10% glycerol, 1 mM TCEP, 5mM MgCl_2_. The clean fractions were concentrated in a 30 kDa cut off concentrator and the protein was flash frozen and stored at −80 °C.

Human Aurora A kinase domain 122-403 and mutants in an N-terminal His-tagged vector (pET30TEV) were transformed into RIL cells alongside the pCDF vector encoding lambda phosphatase. The bacteria were grown in LB at 37 °C until the O.D. at 600 nm reached 0.6-0.8. Expression was then induced with 0.5 mM IPTG overnight at 20 °C. The pelleted cells were resuspended in 10 ml of ice-cold lysis buffer per litre of culture (50 mM TRIS pH 7.5, 250 mM NaCl, 20 mM imidazole, 10% glycerol, 5 mM MgCl_2_, one EDTA-free protease inhibitor tablet per 50 ml of buffer). The resuspended cells were sonicated at 60% amplitude for 10 sec on, 20 sec off, 5 min total to lyse them. The soluble fraction was collected at 17000 rpm for 5 min. After filtering through a 0.45 μm filter the soluble was loaded onto a HisTrap HP column, washed and eluted in lysis buffer using a gradient of maximum 500 mM imidazole. The His-tag was then cleaved overnight using TEV protease in dialysis at 4 °C into 50 mM TRIS pH 7.5, 250 mM NaCl, 10% glycerol, 5 mM MgCl_2_, 1 mM TCEP. After dialysis the cleaved protein was rebound to the HisTrap equilibrated in dialysis buffer and a gradient of 500 mM imidazole was used to elute the tag-free protein. The Aurora A-containing fractions were concentrated in a 10 kDa cut-off concentrator and loaded onto a SD200 16/600 size exclusion column equilibrated into 50 mM TRIS pH 7.5, 200 mM NaCl, 10% glycerol, 5 mM MgCl_2_, 1 mM TCEP. In the final step Aurora A was concentrated again and flash-frozen before storage at −80 °C.

PLK1 binding DARPin in an N-terminal His-tagged vector were transformed into B834 RIL cells. The bacteria were grown in LB at 37 °C until the O.D. at 600 nm reached 0.6-0.8. Expression was then induced with 0.5 mM IPTG overnight at 20 °C. The pelleted cells were resuspended in 10 ml of ice-cold lysis buffer per litre of culture (50 mM TRIS pH 7.5, 500 mM NaCl, 20 mM imidazole, 10% glycerol, 5 mM MgCl_2_, one EDTA-free protease inhibitor tablet per 50ml of buffer). The resuspended cells were lysed by sonication at 60% Amplitude for 10 sec on, 20 sec off, 5 min total. The soluble fraction was collected at 17000 rpm for 5 min. After filtering it through a 0.45 μm filter the soluble fraction was loaded on a HisTrap HP column. The DARPin was eluted in lysis buffer with a gradient of imidazole up to 500 mM maximum, concentrated to under 5 ml in a 5 kDa cut-off concentrator and loaded onto a SD200 16/600 column equilibrated into 50 mM TRIS pH 7.5, 250 mM NaCl, 10% glycerol, 5 mM MgCl_2_, 1 mM TCEP. In the final step the DARPin was concentrated again and flash-frozen before storage at −80 °C.

CEP192 442-533 for use in competition NMR was expressed and purified as in detailed in previous work (Holder *et al*, 2024). TPX2 1-43 for use in competition NMR was expressed and purified as detailed in previous work (McIntyre *et al*, 2017a).

### NMR

For NMR studies Bora 1-120, Bora 18-120 S27A,S41A,T52A and Bora 1-224 were expressed in BL21 (DE3) *E. coli* in 250 mL of ^15^N/^13^C minimal media as His-SUMO-Bora fusions (Holder *et al*, 2024)(Rejnowicz *et al*, 2024). Minimal media contained 2 g/L ^15^NH_4_Cl and 4 g/L ^13^C D-glucose in 50 mM Na_2_HPO_4_, 25 mM KH_2_PO_4_, 20 mM NaCl, supplemented with 2 mM MgSO_4_, 0.2 mM CaCl_2_, 0.01 mM FeSO_4_, a micronutrient cocktail and a vitamin solution (BME vitamins 100× solution, Sigma– Aldrich). Proteins were purified from cell lysate using His-tag affinity chromatography in Tris buffers (50 mM tris, 250/350 mM NaCl, 20 to 500 mM imidazole, 10% glycerol, pH 7.5). The His-SUMO tag was cleaved with TEV and separated by rebinding to the HisTrap column. BORA proteins were polished using size exclusion chromatography into ‘NMR buffer’: 20 mM (K/H)_3_PO_4_ 150 mM NaCl, 2.5% glycerol, pH 6.8. TEV cleavage leaves a short ‘hangover’ sequence GS at the N-terminus.

The following NMR spectra were recorded at 10 °C with a 0.3 mM sample of ^15^N/^13^C-labelled Bora 1-120: ^1^H–^15^N HSQC, ^1^H–^13^C HSQC, HNCO, HNCA, HNCoCA (recorded uniformly), and HNcaCO, HNCACB and HNcocaCB (recorded with NUS). These spectra allowed the complete *ab initio* assignment of the ^1^H–^15^N HSQC spectrum and all accessible HN, H, CO, Ca, and Cb resonances were recorded. An HBHAcoNH spectrum in combination with the ^1^H–^13^C HSQC provided assignments for all but a handful of the Ha and Hb protons. Additional ^1^H–^15^N-HSQC spectra were recorded at 15, 20, 25 and 30 °C. Transverse relaxation rates (*R*_2_) and heteronuclear NOEs (hetNOE) for backbone ^15^N nuclei were measured at 10 °C. Recycle delays were 2.5 s for *R*_2_ and 5.0 s for hetNOE experiments. For *R*_2_, twelve relaxation periods were used ranging from 17 to 204 ms; two were duplicated to help with error estimations. Measurements were limited to those residues with independent, resolvable peaks in the ^1^H–^15^N HSQC spectrum. Peak intensities were measured and relaxation rates/hetNOE values analysed using PINT (Ahlner *et al*, 2013).

^1^H–^15^N HSQC assignment for an equivalent 0.3 mM sample of Bora 18-120 S27A,S41A,T52A was achieved through comparison with Bora 1-120 and using a HNCA/HNCoCA pair of spectra to confirm assignments at the N-terminus and close to mutation sites.

### Bora phosphorylation in NMR

Wild-type Aurora-A 122-403 (final conc. 1 µM) was added to a 30 µM sample of ^15^N/^13^C-labelled Bora 1-120 in NMR buffer supplemented with 2 mM ATP and 5 mM MgCl_2_ at 20 °C. Phosphorylation status after addition of Aurora-A was tracked by recording sequential ^1^H–^15^N HSQC spectra. The identity of pSer59 was confirmed using an HNCA experiment at 10 °C, linking through (*i*,*i* – 1) C⍺ resonances for shifted peaks. Clear ^1^H-^15^N chemical shift perturbations (>0.02 ppm) were shown for residues between Thr52 and Ile71 (inclusive).

Similarly, for ERK phosphorylation, ERK1 or ERK2 (final conc. ∼1 µM) was added to 50 uM ^15^N/^13^C-labelled samples of Bora 1-120 or Bora 18-120 S27A,S41A,T52A in NMR buffer supplemented with 1 mM ATP and 3 mM MgCl_2_ at 20 °C. Phosphorylation status after addition of kinase was tracked by recording sequential ^1^H–^15^N HSQC spectra. The identity of the phosphorylated residues (pSer112 for Bora 18-120 S27A,S41A,T52A) was confirmed using an HNCA experiment or HNCA/HNcoCA pair at 10 °C.

### NMR titrations/competition assays

For binding studies, concentrated Aurora-A 122-403 C290A/C393A was added to 30– 80 uM samples of isotopically labelled human Bora 1-120 or Bora 18-120 S27A, S41A, T52A in NMR buffer in small <u>aliquots</u> up to a 1:1 molar ratio. For competition assays, Aurora-A was first added up to a 1:2.5 molar ratio of [AurA]:[Bora] and then TPX2 1-43 or CEP192 442-533 was added in small, concentrated aliquots up to a final <u>molar</u> <u>ratio</u> of 1:2.5:1. Peak intensities were ratioed to the original Bora-only spectrum; the change in Bora concentration was small.

### PLK1 pThr210 phosphorylation assay

To create phosphorylated Bora, Bora WT and mutants were preincubated at 5 μM with 640 nM ERK2 in 50 μl of phosphorylation buffer (25 mM Hepes pH 7.5, 150 mM NaCl, 1 mM DTT, 20 mM MgCl_2_, 200 μM ATP) for 2 h at 30 °C. The phosphorylated Bora at a final concentration of 1 μM was then mixed with 200 nM of PLK1 K82R 3-330 and Aurora-A 122-403 in 30 μl total phosphorylation buffer. This was incubated at 30 °C for 30 min before 30 μl of 2 X SDS loading dye was added, the samples were boiled and were analysed via western blot with transfer onto a nitrocellulose membrane. The blots were blocked in 5% milk in Tris buffered saline with 0.01% Tween 20. The membranes were split and probed with anti-Bora and anti-PLK1 pT210 antibodies. Fluorescent secondary antibodies from Biorad were used and the western blot imaged using the iBright imaging system (Invitrogen). The iBright analysis software was used to quantify the levels of fluorescence using the local background corrected volume. When analysing the effects of S59 phosphorylation in Bora, there was an extra 30 min incubation period of the phosphorylated Bora with Aurora-A before the PLK1 was added.

The levels of all proteins included in the phosphorylation assay were assessed using the SYPRO Ruby staining of the nitrocellulose membrane (SYPRO Ruby protein blot stain, Invitrogen), with visualisation on the iBright imaging system (Invitrogen).

### Modelling of ternary complexes

The Google Colab version of AlphaFold2 multimer in local mode was used to model a three-way complex between human Bora (Q6PGQ7), human PLK1 (P53350) and human Aurora-A (O14965) (Mirdita *et al*, 2022)(Evans *et al*, 2021). 10 seeds were used in the modelling with 6 recycles. The top ipTM score from the 50 models is included in supplementary table 3. Modelling the complexes between Aurora-A kinase, PLK1 and a phosphorylated version of Bora was achieved using the alphafoldserver.com and AlphaFold3 (Abramson *et al*, 2024). The full-length sequences of Aurora-A, Bora and PLK1 were taken from Uniprot.

Models used in this study were deposited to ModelArchive (Tauriello *et al*, 2025). Pairwise ipTM scores were extracted from the json output from alphafoldserver.com. 10 models were generated for each complex, with all 5 ipTM scores extracted from each model and plotted on a violin plot using Origin (OriginPro version 2024). AlphaBridge was used to identify the predicted interfaces between proteins complexes modelled on AlphaFold3 without ADP or phosphorylation present (Álvarez-Salmoral *et al*, 2024).

### Phylogenetic analysis

Orthologous sequences were identified using iterative PSI-BLAST searches on the non-redundant database using the UniProt sequence Q6PGQ7 Bora 1-559, P53350 PLK1 1-603 or O14965 Aurora-A 1-403 as a template. Diverse orthologues were identified by constraining the searches to use just a particular taxonomic group. The multiple sequence alignments of PLK1 and Bora were generated using MAFFT (Katoh *et al*, 2017). Sequence conservation logos were created using the MAFFT alignment and WebLogo (Crooks *et al*, 2004). The *Strongylocentrotus purpuratus* orthologues used in the AlphaFold3 modelling were XP_030849245.1 (PLK1, 1-330), XP_030831330.1 (Aurora-A 80-346) and XP_030848668.1 (Bora 18-112).

### Analytical SEC

A superose 12 10/300 column (Cytiva) was equilibrated in 50 mM tris pH 7.5, 250 mM NaCl, 10% glycerol, 5 mM magnesium chloride, 0.5 mM TCEP. PLK1 3-330 K82R or K82R R106A S99R and DARPin were mixed at 1:1 molar ratio in a total volume of 500 μl. The mixed sample and the individual proteins were subject to SEC in sequential runs and the resulting traces overlaid.

### Fluorescence anisotropy-based assays

Assays were performed in 25 mM HEPES pH 7.5, 150 mM NaCl, 2 mM DTT, 5 mM MgCl_2_, 0.01% Tween 20. For CEP192 and TPX2 competition assays 10 μM Aurora-A kinase domain (unphosphorylated C290A C393A 122–403) was preincubated on ice with 100 nM FAM-TPX2 7-43 or 100 nM FAM-CEP192 501–533. Serial dilutions of competitor (Bora) were performed in triplicate into a low volume 384-well black plate, before the premixed Aurora-A and tracer were added. A control row included no tracer. Plates were left incubating for 1 h at room temperature before the fluorescence anisotropy was measured using a HIDEX plate reader, with excitation at 485 nm and emission at 535 nm.

For direct binding experiments, the assay was performed as above, but PLK1 (3-330 K82R or K82R R106A S99R) was diluted across the plate from high to low concentration in 25 mM HEPES pH 7.5, 50 mM NaCl, 2 mM DTT, 5 mM MgCl_2_, 0.01% Tween 20, before the addition of 50 nM tracer (FAM-Bora 52-73 wild-type) to three replicate rows, whereas only buffer was added to the control row.

Fluorescence anisotropy data were processed using Microsoft Excel to calculate intensity and anisotropy using the equation listed in (Yeo *et al*, 2013). Data was fitted on Origin (OriginPro version 2024).

### Peptide synthesis

Peptide synthesis was performed using Liberty Blue peptide synthesiser (CEM Corporation) with microwave heating at 0.1 mmol scale on Rink Amide ProTide resin (loading 0.19 mmol/g). Standard preprogramed coupling and deprotection cycles were applied. The deprotection was achieved using 20% piperidine in DMF with microwave heating at 90 °C for 100 s, followed by three DMF washing steps. The couplings were performed using 5 eq. of Fmoc-protected amino acid, 5 eq. of N,N’-diisopropylcarbodiimide (DIC) and 5 eq. of 2-cyano-2-(hydroxyimino)acetate (Oxyma) in DMF with microwave heating at 90 °C for 3 min followed by two DMF washing steps. Single couplings were applied to the first 15 residues while two coupling cycles were used for residue 16 and further.

The resin with synthesized peptide was then transferred to SPS tube and labelled using 3 eq. of 5(6)-carboxyfluorescein, 3 eq. of DIC and 3 eq. of Oxyma in DMF for 16h, followed by washing with 10 ml of 20% piperidine in DMF two times for 5 min and three times with 10 ml DMF. After further washing three times with 10 ml dichloromethane and two times with 10 ml of diethylether the resin was dried under vacuum for 30 min. To deprotect the side chains and cleave the peptide from the resin, the resin was incubated on a rotator for 3 h with 10 ml of cleavage mix (92.5% trifluoroacetic acid (TFA), 2.5% water, 2.5% triisopropylsilane (TIPS), 2.5% 3,6-dioxa-1,8-octanedithiol (DODT)) and filtered. The filtrate was concentrated to ca 1 ml under a stream of nitrogen and the peptide was precipitated by addition of 10 ml of ice-cold diethylether and isolated by centrifugation (6000 rpm for 5 min). The precipitate was resuspended in 10 ml of ice-cold diethylether and isolated by repeating the centrifugation step. After decanting the diethylether, the precipitate was allowed to dry for 30 min, dissolved in 5 ml of 1% (v/v) acetic acid and freeze dried.

### Peptide purification

Peptides were dissolved in 4-10 ml of 1:1 mixture of acetonitrile and water and purified using Agilent 1260 infinity system equipped with UV detector and fraction collector on Kinetex EVO 5 µm C18 100Å 21.2 x 250 mm reverse phase column. 1-4 ml of the peptide solution was injected and 25 min gradient of 20-40% acetonitrile in water with 0.1% formic acid additive was run at 10 ml/min. The fractions containing peptide were pooled and freeze dried.

The identity of the peptides was confirmed by high-resolution mass spectrometry on Bruker Maxis Impact spectrometer using electrospray ionisation. The purity was determined by analytical HPLC on Agilent 1290 Infinity II system using Ascentis peptide column and gradient 5-95% of acetonitrile in water with 0.1% trifluoroacetic acid additive at 0.5 ml/min for 10 min.

### Mass spectrometry analysis

Bora 18-120 wild-type and mutants were diluted to 8 µM in 25 mM Hepes pH 7.5, 150 mM NaCl, 20 mM MgCl_2_, 1 mM DTT and 200 µM ATP with and without inclusion of 1.5 µM ERK2. After incubation at 30 °C for 2 hours, EDTA was added to a final concentration of 20 mM to stop any further phosphorylation. The samples were flash frozen and stored ahead of analysis.

Accurate mass spectra were acquired on an Impact II QqTOF spectrometer equipped with a VIPHESI source using either electrospray or atmospheric pressure chemical ionisation. Samples were introduced using an HTC PAL autosampler and Bruker Elute Pump. HPLC columns were heated to 40 °C unless otherwise stated. Samples passed through a Bruker Diode array uv-detector before entering the mass spectrometer. Calibration was performed by infusion of 5 mM sodium formate solution at the end of each acquisition. The guard column used was Waters Acquity Vanguard Protein BEH C4 300A 1.7µm 2.1mm x 5mm (p/n 186004623), with the column being Waters Acquity Vanguard Protein BEH C4 300A 1.7µm 2.1mm x 100mm (p/n 186004496). The separation was undertaken in water and acetonitrile, both containing 0.1% formic acid. Maximum entropy deconvolution methods were used as part of the processing.

### Peptide array

Peptide arrays covering 15 amino acid peptides of human Bora 1-559 with a 5 residue shift were synthesized on 10×15 cm cellulose membrane made of 6-aminohexanoic acid modified Whatman 540 filter paper, using MultiPep 2 peptide synthesizer (CEM Corporation). First, Fmoc-8-amino-3,6-dioxaoctanoic acid spacer was coupled to the membrane to provide a distance between the solid support and the peptide. Peptides were then assembled using standard Fmoc-based solid phase synthesis with double deprotection step (2 x 15 min) using 20% piperidine in DMF, double couplings (2 x 30 min) with DIC and Oxyma as coupling reagents and 10% acetic anhydride in DMF as capping reagent (15 min). Final deprotection was achieved by incubating the membrane in the mixture of 92.5% TFA, 2.5% water, 2.5% TIPS and 2.5% DODT for 3 h, followed by three dichloromethane, three DMF, three ethanol washes and air-drying.

The peptides on the membrane were rehydrated in ethanol for 5 min followed by 5 min wash in TBST (50 mM Tris pH 7.5, 150 mM NaCl, 0.01% Tween-20). The membrane was blocked with 1% bovine serum albumin (BSA) in binding buffer (25 mM TRIS pH 7.5, 150 mM NaCl, 5 mM MgCl_2_, 1 mM DTT, 0.01% Tween-20, 10% Glycerol) with agitation for 1 h and washed for 5 min in binding buffer. The membrane was then incubated with 100 nM of autophosphorylated Aurora-A 122-403 and 100 µM ATP in binding buffer for 1 h for phosphorylation to occur. The protein was drained off and the membrane washed using 2 x 5 min wash of 10% SDS and 5 x 5 min wash in deionised water. The water was drained off and the membrane incubated in Pro-Q Diamond Phosphoprotein stain (ThermoFisher Scientific) for 1 h. The stain was removed and the membrane washed in 2 x 5 min washes with deionised water. The membrane was detained several times in 50 mM NaOAc pH 4.0 + 5% Acetonitrile. The level of phosphorylation was then imaged using a Gel Doc system.

## Author contributions statement

JAM – Conceptualization. Writing - original draft. Investigation. Formal analysis. Visualization. Validation. Methodology. Writing—review and editing.

MB - Writing - original draft. Investigation. Formal analysis. Visualization. Methodology. Writing—review and editing.

MW – Writing - original draft. Resources. Investigation. Formal analysis. Visualization.

VG – Investigation.

AJW- Conceptualization. Funding acquisition.

MHW - Conceptualization. Funding acquisition. Supervision.

RB – Conceptualization. Funding acquisition. Supervision. Writing – original draft. Writing—review and editing.

## Acknowledgements

This research was supported by the Biotechnology and Biological Sciences Research Council (BBSRC) (BB/V003577/1 and BB/V003577/2). We thank Arnout Kalverda for NMR support and Roland Dunbrack Jr (Fox Chase Cancer Centre) for help with modelling. Facilities for NMR spectroscopy were funded by the University of Leeds (ABSL award) and Wellcome Trust (108466/Z/15/Z).

## Additional information

The assignment for Bora 1-120 has been deposited at the BMRB under code 52970.

The ternary complex models of the human orthologues were deposited to the ModelArchive under codes ma-1klkx (PLK1 21-330, Bora 18-120 with Aurora-A 122-403 with AlphaFold3), ma-uptw5 (Full-length PLK1, Bora and Aurora-A with AlphaFold2), ma-x9q8r (PLK1 21-330, Bora 18-120 with Aurora-A 122-403 with AlphaFold2), ma-bxipa (Full-length PLK1, Bora and Aurora-A with AlphaFold3) and ma-bldfw (PLK1 21-330, Bora 18-120 with Aurora-A 122-403 with AlphaFold3 including phosphorylation of Bora S59 and S112).

