## Supplementary Information for "Bora Bridges Aurora-A Activation and Substrate Recognition of PLK1"

### List of contents

Figures

Figure S1 – Modelling the ternary complex between Bora, PLK1 and Aurora-A

Figure S2 – Comparing the complex models

Figure S3 – Modelling the binary complex between Bora and PLK1

Figure S4 – Biochemical analysis of mutations in the Bora:PLK1 interface

Figure S5 – Bora sequence alignment

Figure S6 – PLK1 kinase domain sequence alignment

Figure S7 – Full assignment of Bora 1-120

Figure S8 – NMR characterisation of Bora 1-120

Figure S9 – Phosphorylation of Bora 1-120 by Aurora-A and ERK2

Figure S10 – Comparison of the predicted Bora:Aurora-A structure with INCENP bound to Aurora-C

Figure S11 – Mass spec analysis of Bora variants in the absence and presence of ERK2.

Tables

Table S1 – Primers used for site directed mutagenesis in this study

Table S2 – Constructs used in this study

Table S3 – iPTM scores from AlphaFold2 and AlphaFold3 predictions

Table S4 – Mass spectrometry analysis of modified Bora samples

Fig S1.

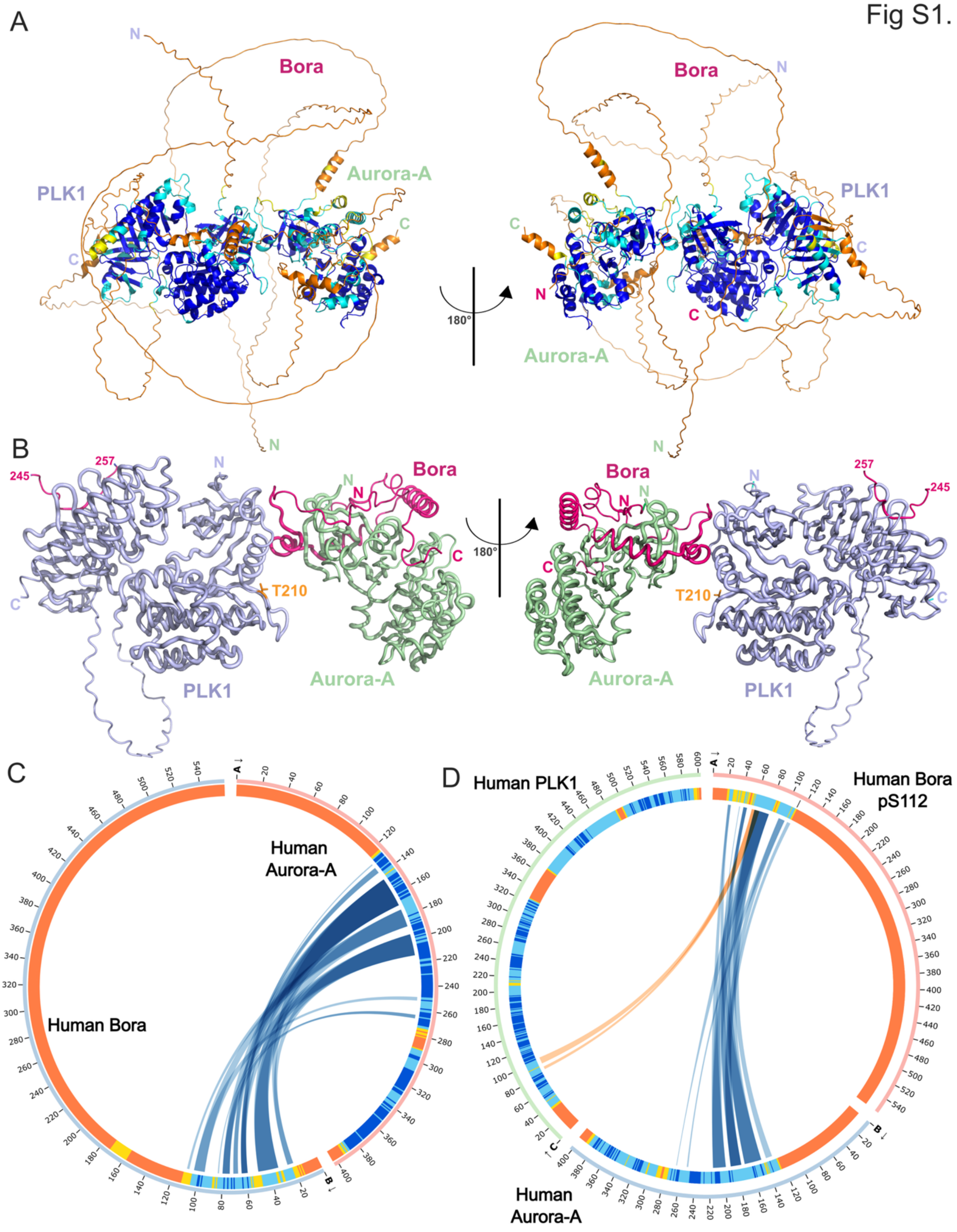
A
Bora
PLK1
Aurora-A
N
C
180°
B
Bora
PLK1
Aurora-A
245
257
T210
180°
C
Human Aurora-A
Human Bora
D
Human PLK1
Human Bora pS112
Human Aurora-A

**Figure S1 - Modelling the ternary complex between Bora, PLK1 and Aurora-A**

**A:** Model of the complex of full-length human Bora bound to full-length Aurora-A and full-length PLK1 from AlphaFold3. The model is coloured according to the pLDDT scores of each residue in the modelling, with most of the model containing intrinsically disordered regions of low confidence. The blue regions have the highest confidence, and the orange regions have the lowest confidence.

**B:** Simplified model removing the low confidence disordered regions, with Bora shown in magenta, Aurora-A shown in green and PLK1 shown in blue. The short Bora peptide covering 245-257 is modelled interacting with the polo-box domain of PLK1.

**C:** AlphaBridge summary of the predicted interfaces between full-length human Aurora-A and Bora from an AlphaFold3 model. The colouring is based on the pLDDT score.

**D:** AlphaBridge summary of the predicted interfaces between full-length Aurora-A, PLK1 and pS112 Bora from an AlphaFold3 model. The colouring is based on the pLDDT score.

Fig. S2

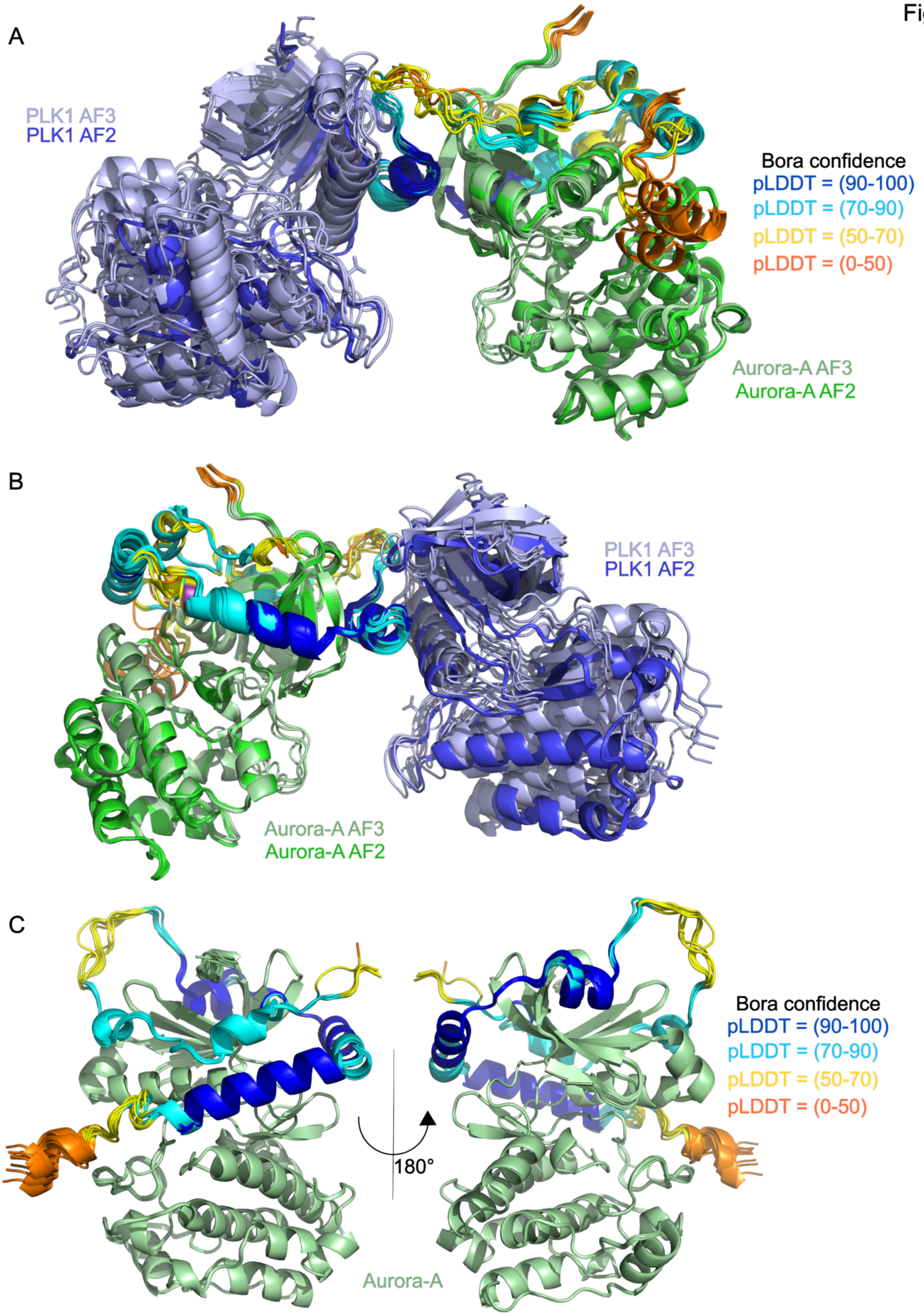


**Figure S2– Comparing the complex models**

**A:** Model of the complex of human Bora 18-120 bound to Aurora-A 122-403 and PLK1 30-330 from both AlphaFold2 and AlphaFold3. The Bora in each model is coloured according to the pLDDT scores of each residue, with most of the model containing intrinsically disordered regions of low confidence. Aurora-A models from AlphaFold3 are shown in light blue, AlphaFold2 models in blue. PLK1 models from AlphaFold3 are shown in light green, AlphaFold2 models in bright green. The 5 top ranked models from each version of AlphaFold are shown, with the models aligned on the Aurora-A kinase domain.

**B:** 180 degree rotation of the model shown in Figure S2A.

**C:** Model of the complex of human Bora 18-120 bound to Aurora-A 122-403 from AlphaFold3. The Bora in each model is coloured according to the pLDDT scores of each residue. Aurora-A is shown in light blue. 10 models have been overlaid, indicating that there is a high degree of consistency between the models.

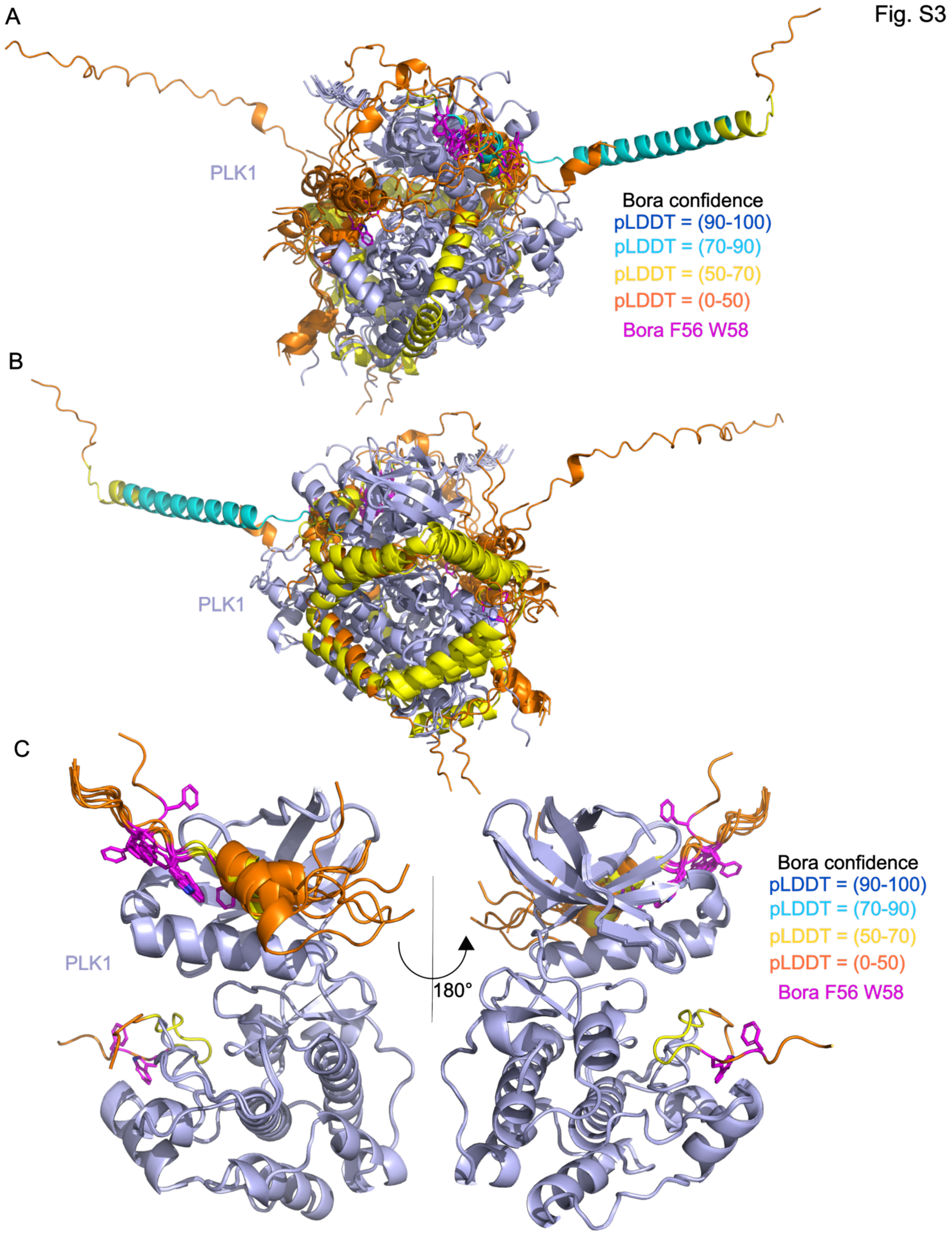


**Figure S3 - Modelling the binary complex between Bora and PLK1**

**A:** Model of the complex of human Bora 18-120 bound to PLK1 30-330 from AlphaFold3. The Bora in each model is coloured according to the pLDDT scores of each residue, with most of the model containing intrinsically disordered regions of low confidence. PLK1 is shown in light blue. 10 models have been overlaid, indicating that there is high variability between them. The two hydrophobic residues that are predicted to interact with PLK1 are shown in pink (Phe56 and Trp58).

**B:** 180 degree rotation of the model shown in Figure S3A.

**C:** Model of the complex of human Bora 52-72 bound to PLK1 30-330 from AlphaFold3. The Bora in each model is coloured according to the pLDDT scores of each residue, with most of the model containing intrinsically disordered regions of low confidence. PLK1 is shown in light blue. The two hydrophobic residues that are predicted to interact with PLK1 are shown in pink (Phe56 and Trp58). 10 models have been overlaid, with all but one placing the two hydrophobic residues in the 'FW' pocket in PLK1.

Fig. S4

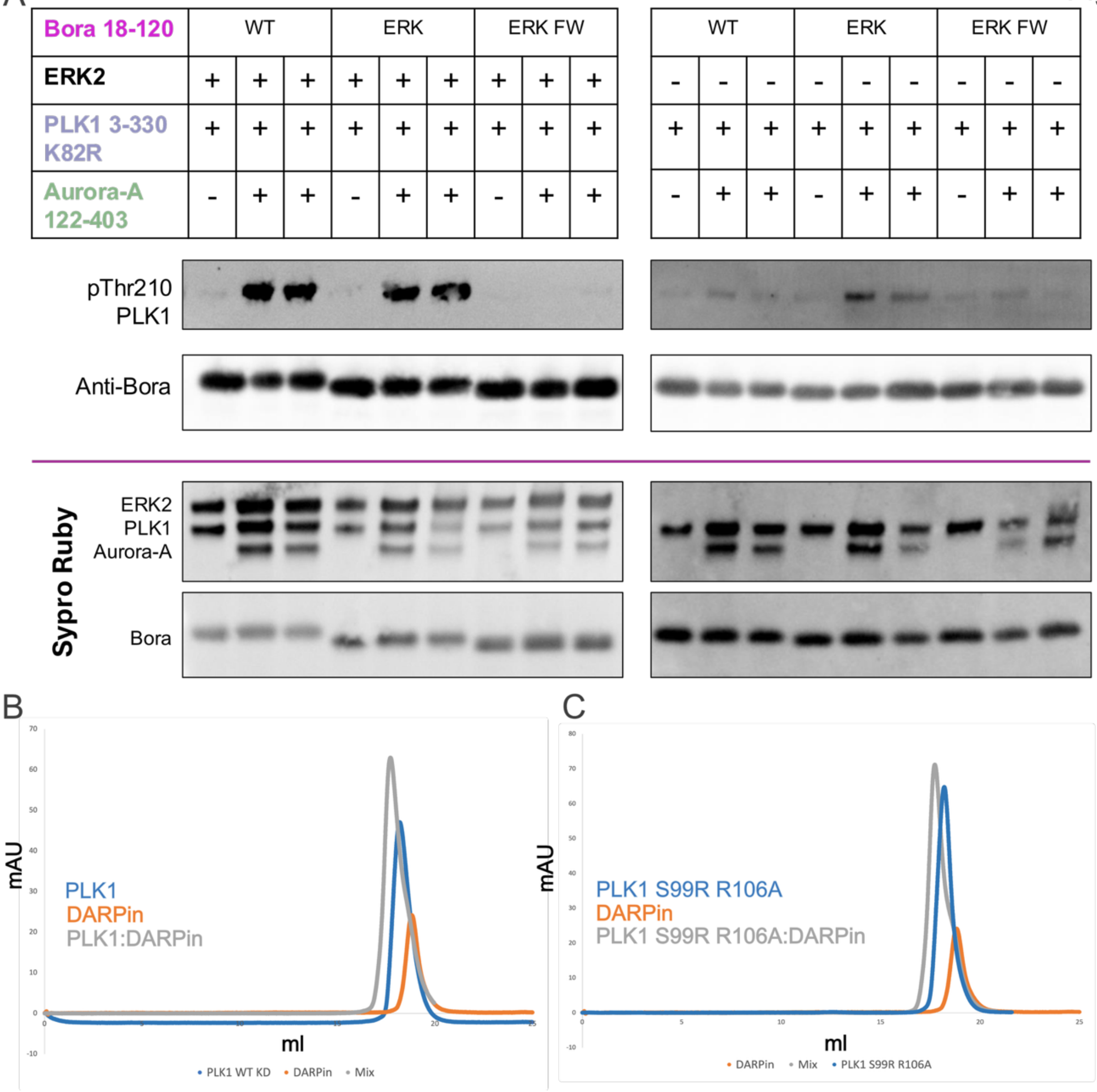


**Figure S4 – Biochemical analysis of mutations in the Bora:PLK1 interface**

**A:** Western blot analysis of the levels of phosphorylation of PLK1 kinase domain at Thr210 when Bora wild-type and mutants (ERK = S27A S41A T52A, ERK FW = S27A S41A T52A F56A W58A) are preincubated with ERK2 and added to wild-type dephosphorylated Aurora-A kinase domain. Mutating the other potential sites phosphorylated by ERK2 does not affect how well Bora can stimulate the activity of Aurora-A (lanes 2 and 3 compared to lanes 5 and 6). When the predicted PLK1 binding sites are also mutated this significantly reduces the level of PLK1 phosphorylation observed (lanes 5 and 6 compared to 8 and 9).

**B:** Analytical SEC analysis of PLK1 K82R 3-330 interacting with the PLK1 binding DARPin. The trace from PLK1 alone is shown in blue, with the DARPin alone in orange and the complex shown in grey.

**C**: Analytical SEC analysis of PLK1 K82R S99R R106A 30-330 interacting with the PLK1 binding DARPin. The trace from PLK1 K82R S99R R106A alone is shown in blue, with the DARPin alone in orange and the complex shown in grey.

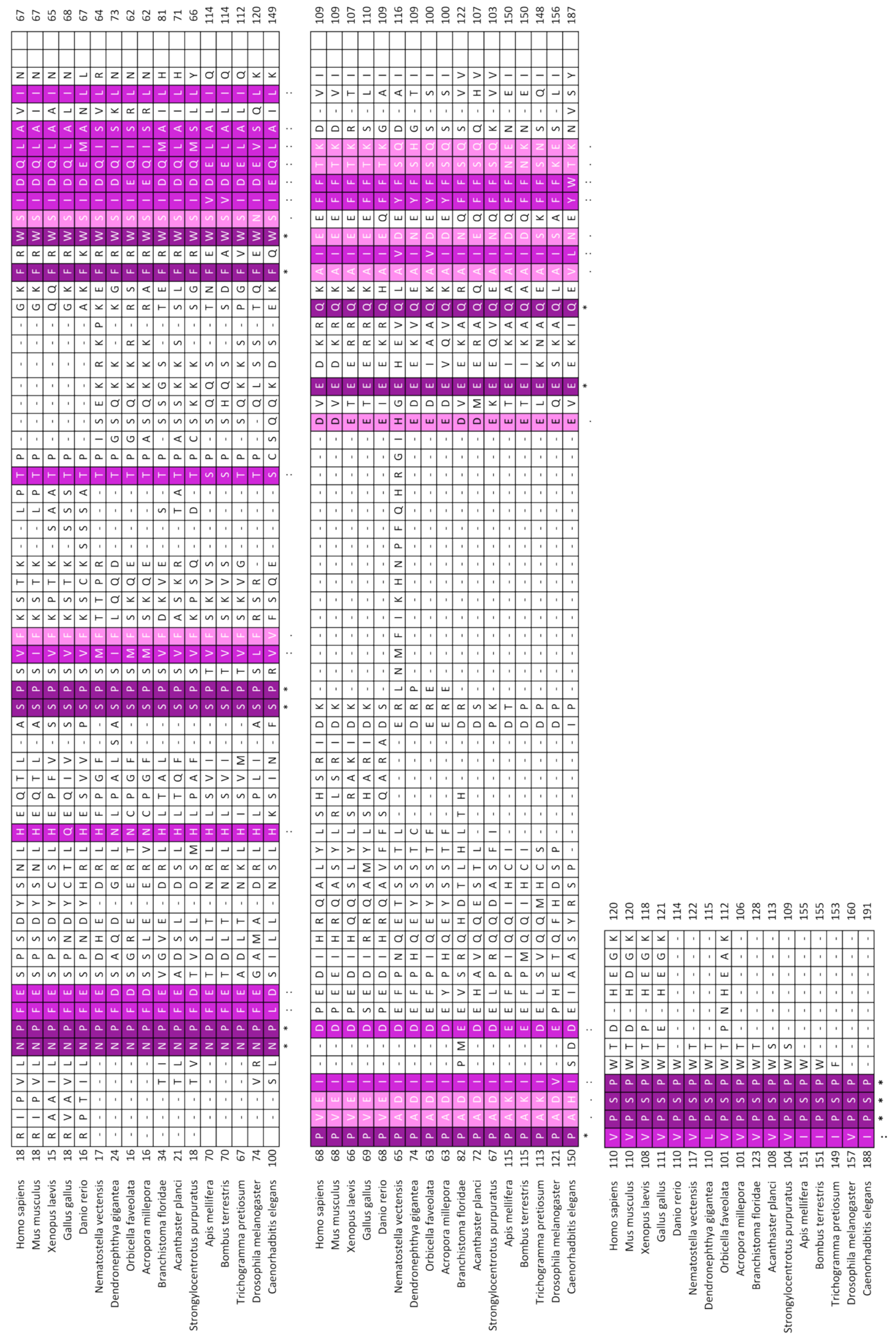


**Figure S5 – Bora sequence alignment**

Sequence alignment of Bora orthologues using MAFFT. Only the conservation over residues 18-120 in *Homo sapiens* Bora is shown. The alignment is coloured according to conservation, with the darker residues being the most conserved.

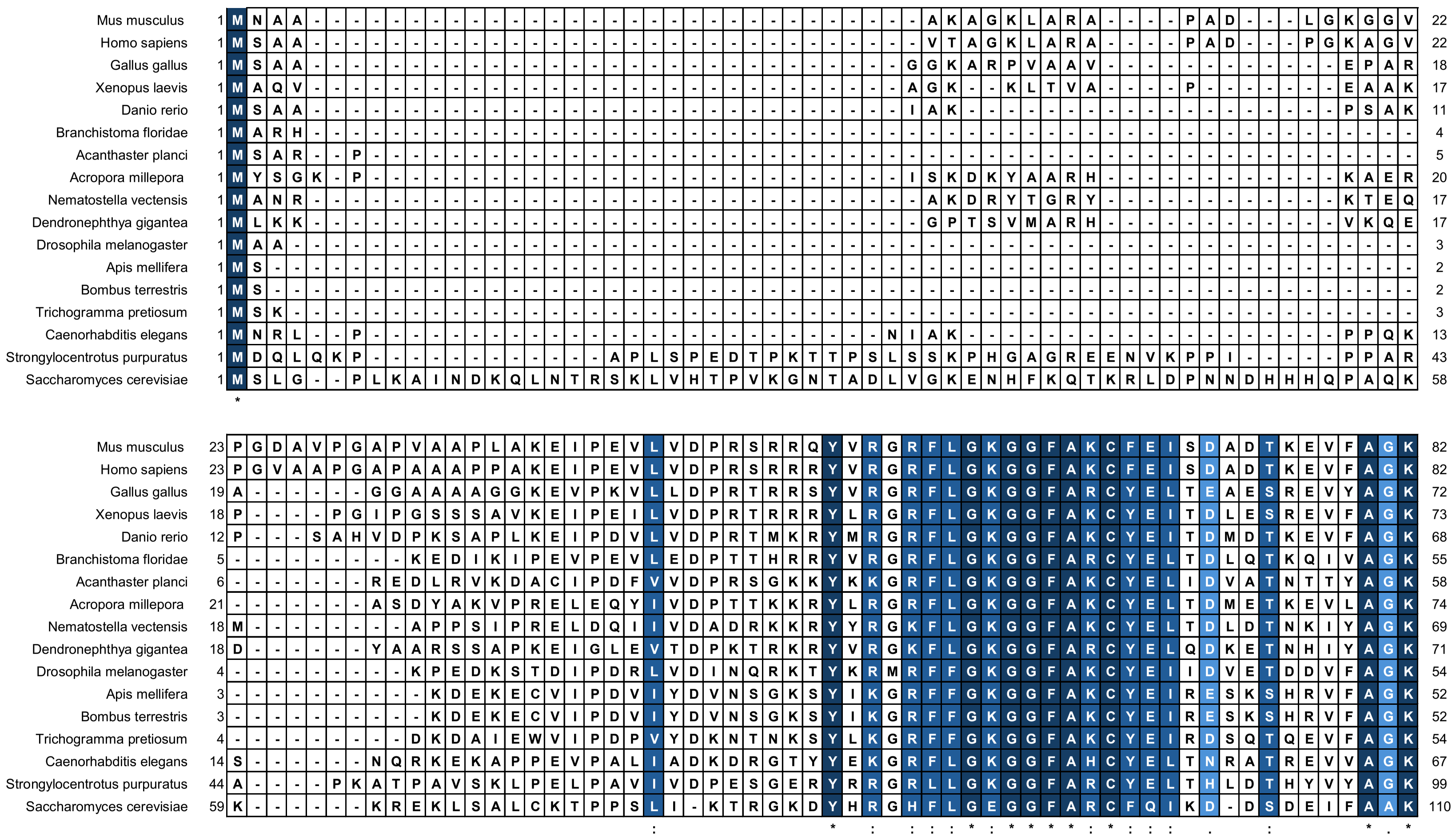
Mus musculus
Homo sapiens
Gallus gallus
Xenopus laevis
Danio rerio
Branchistoma floridae
Acanthaster planci
Acropora millepora
Nematostella vectensis
Dendronephthya gigantea
Drosophila melanogaster
Apis mellifera
Bombus terrestris
Trichogramma pretiosum
Caenorhabditis elegans
Strongylocentrotus purpuratus
Saccharomyces cerevisiae

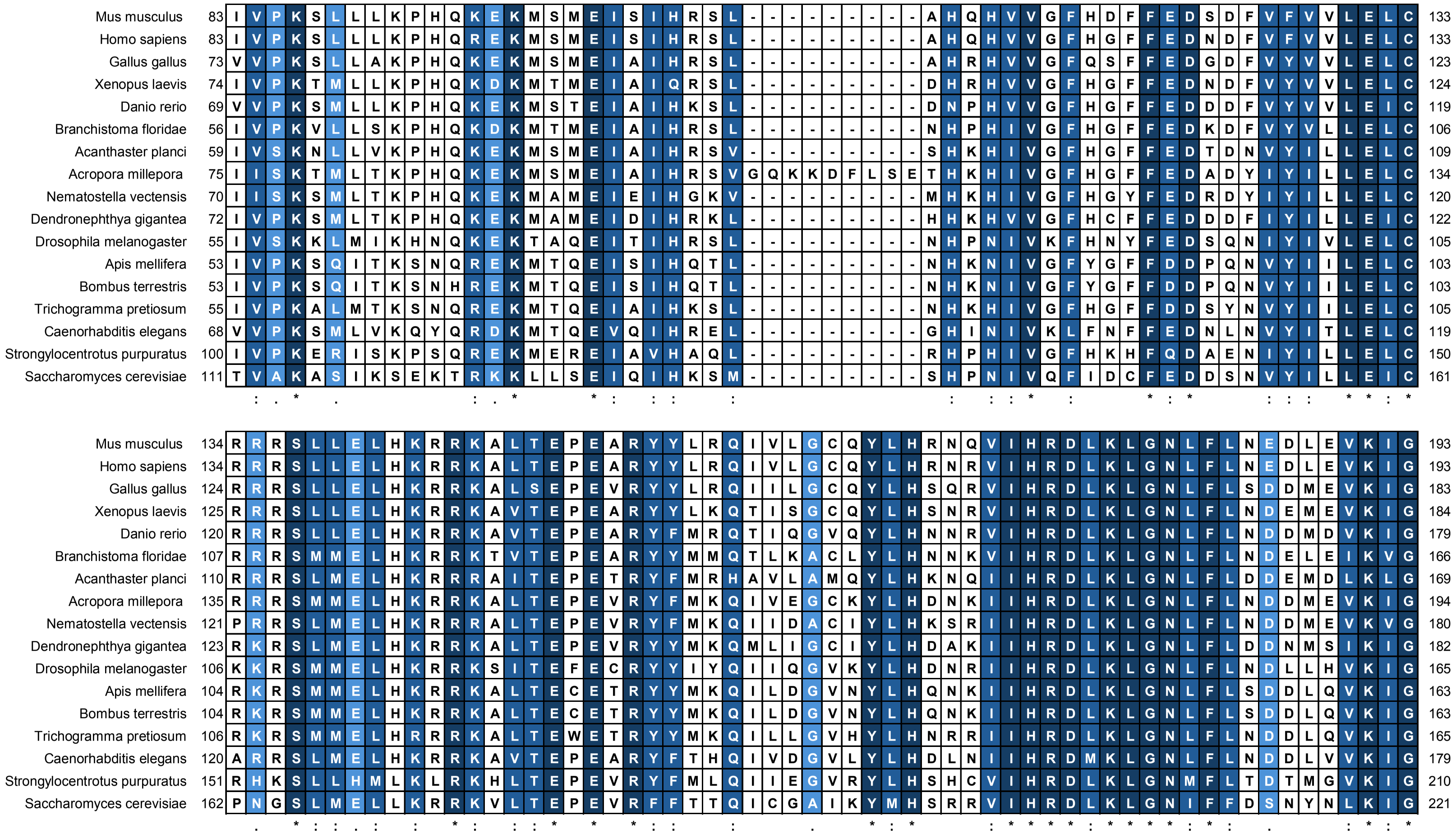

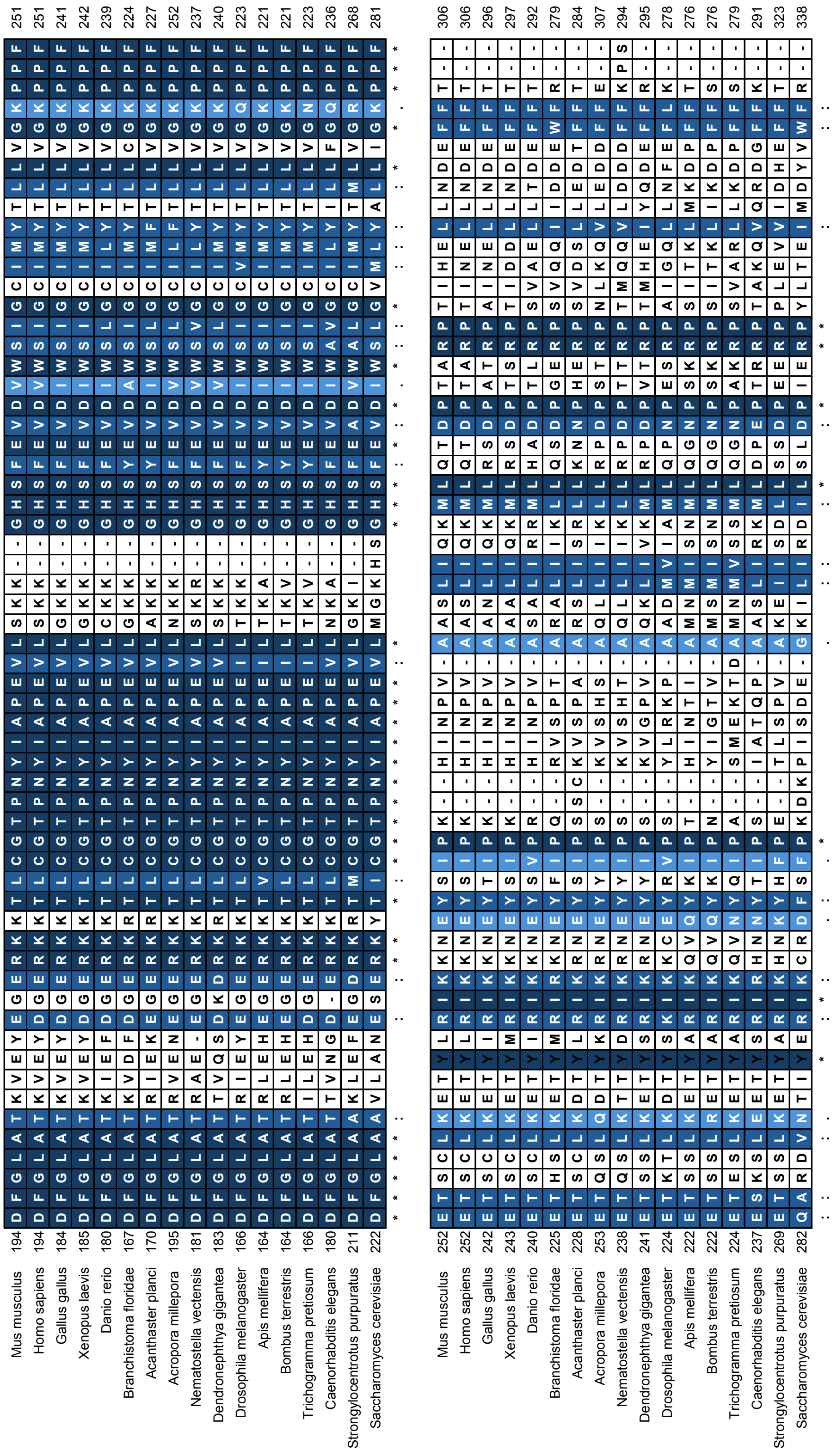
Mus musculus
Homo sapiens
Gallus gallus
Xenopus laevis
Danio rerio
Branchistoma floridae
Acanthaster planci
Acropora millepora
Nematostella vectensis
Dendronephthya gigantea
Drosophila melanogaster
Apis mellifera
Bombus terrestris
Trichogramma pretiosum
Caenorhabditis elegans
Strongylocentrotus purpuratus
Saccharomyces cerevisiae

```
Mus musculus                  307 SGYIPARL-PITCLTIPPRFSIAPSS---------------------LDPS--------  335
Homo sapiens                  307 SGYIPARL-PITCLTIPPRFSIAPSS---------------------LDPS--------- 335
Gallus gallus                 297 SGYIPSRL-PTSCLTVAPRFSIAPST---------------------LELN--------- 325
Xenopus laevis                298 SGYIPSRL-PTTCLTVPPRFSIAPST---------------------IDQS--------- 326
Danio rerio                   293 SGYAPMRL-PTSCLTVPPRFSIAPSS---------------------LDPAL-------- 322
Branchistoma floridae         280 AGYMPSRL-PVTCLTMAPRFPASAT----------------------------------- 303
Acanthaster planci            285 SDYLPPKL-PTTCLTVTPRFAPLVT----------------------------------- 308
Acropora millepora            308 -GFTPNHL-PISSLTMAPRFATTSSAN-------------------LLGL---------- 336
Nematostella vectensis        295 SGYMPTRL-PVSCLTMPPRFATGSSSN-------------------LLQPG--------- 325
Dendronephthya gigantea       296 SGYLPSTL-PASALSMAPKFNPSTSI---------------------LEPKAIGSGMMQK 333
Drosophila melanogaster       279 GSKVPMFL-PSSCLTMAPRIGSNDT----------------------IEDSM-------- 307
Apis mellifera                277 CGFMPLSL-PLSCLTMAPRLDM-------------------------LEMHN-------- 302
Bombus terrestris             277 SGFMPHSL-PLSCLTMAPRLDL-------------------------LEMHN-------- 302
Trichogramma pretiosum        280 HGFMPASL-PSSCLTMAPRLDM-------------------------LEAL--------- 304
Caenorhabditis elegans        292 SGFMPTRL-PVSCLTMVPKFGGHETSM--------------------MEENVAPRGVDAR 330
Strongylocentrotus purpuratus 324 QGFVPETLSPTACVT-APDFPITKVLISDRKGDLVTNEFCRLKRESTLQRRD-------- 374
Saccharomyces cerevisiae      339 -GTFPPSI-PSTVMSEAPNFEDIPEEQS-----LVNFKDC-MEKSLLLESMSSDKIQRQK 390
                                   .  *  : * : ::  * :
```

**Figure S6 – PLK1 kinase domain sequence alignment**

Sequence alignment of PLK1 orthologues using MAFFT. Only the alignment of the kinase domain is shown. The alignment is coloured according to conservation, with the darker residues being the most conserved.

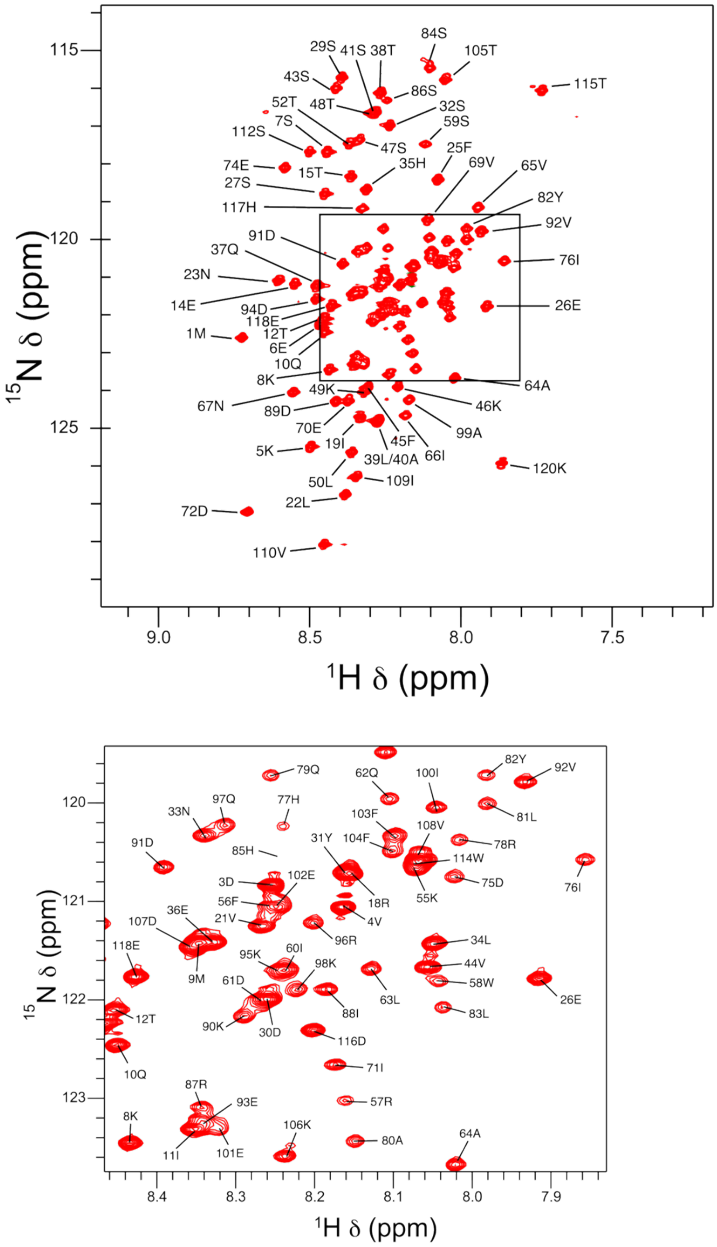


**Figure S7: Full assignment of Bora 1-120**

Assigned $^{1}$H-$^{15}$N core HSQC spectrum of Bora 1-120 at 10 °C. The Gly backbone NH and Trp/Gln/Asn/Arg side-chain regions are omitted to simplify the visualisation. The panel below shows an expanded version of the congested central region.

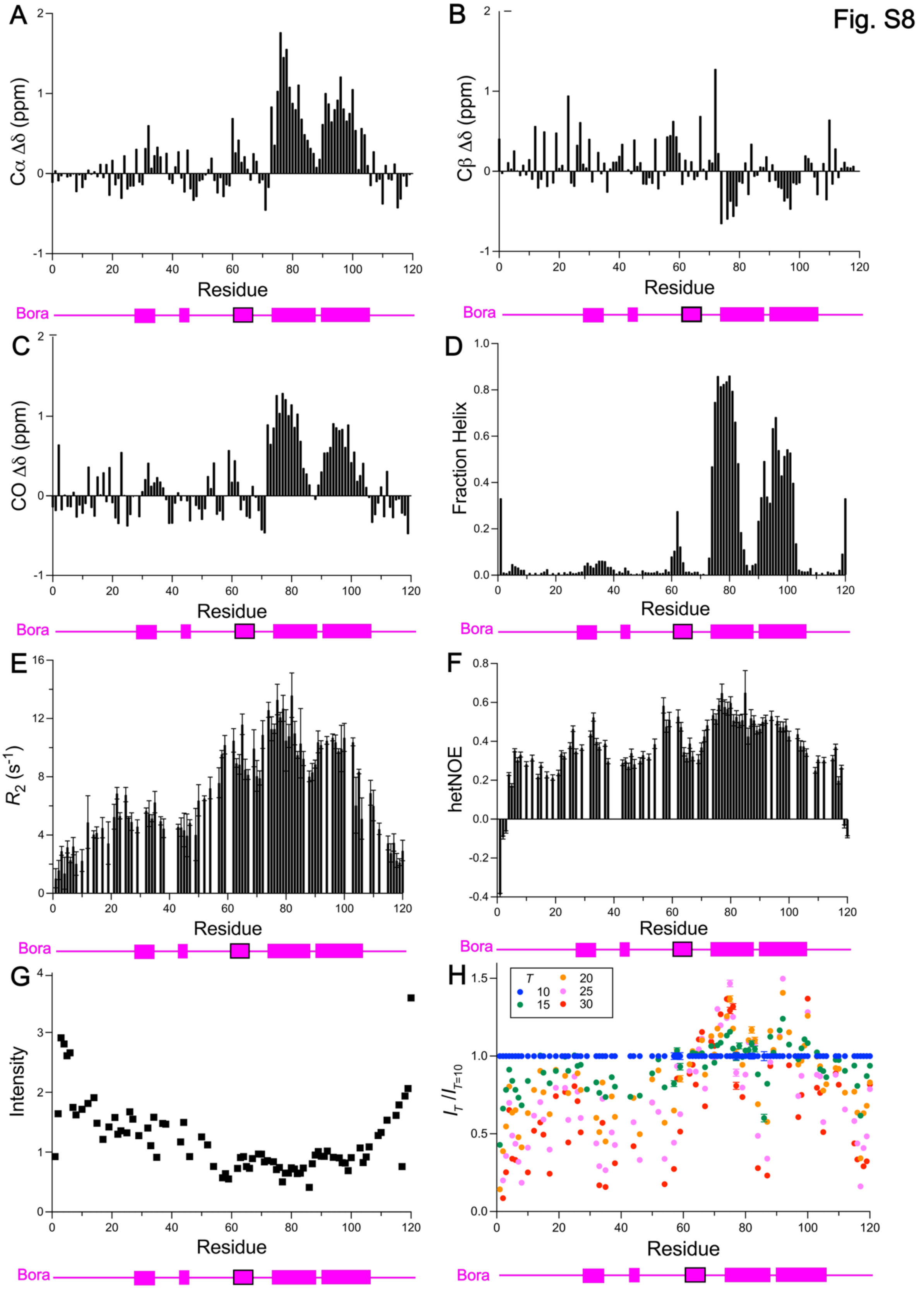


**Figure S8 - NMR characterisation of Bora 1-120**

**A-C:** Secondary shifts ($\Delta\delta$) for C$\alpha$, (**B**), CO (**C**) and C$\beta$ (**D**) nuclei in Bora 1-120. Reference coil values were generated using a web server hosted by the University of Copenhagen (https://www1.bio.ku.dk/english/research/bms/sbinlab/randomchemicalshifts2/[1[1]). Coil values are subtracted from measured shift values to give secondary shifts ($\Delta\delta = \delta - \delta_{ref}$). A representation of the model-predicted Bora secondary structure (in complex with Aurora-A or Aurora-A/PLK1) is shown below in pink, with helices represented as rectangles. The helix suggested to interact with the PLK1 'FW' pocket is bordered in black.

**D:** Residue specific helical propensities within Bora 1-120 calculated using a TALOS-N analysis of all backbone chemical shifts [2]. The analysis indicates two regions of high helical propensity towards the C-terminus. A representation of the model-predicted Bora secondary structure is shown below in pink, with helices represented as rectangles. The helix suggested to interact with the PLK1 'FW' pocket is bordered in black.

**E:** Transverse relaxation rates ($R_2$) for backbone $^{15}$N nuclei in Bora 1-120. A representation of the model-predicted Bora secondary structure is shown below in pink, with helices represented as rectangles. The helix suggested to interact with the PLK1 'FW' pocket is bordered in black.

**F:** Heteronuclear NOE values for HN groups in Bora 1-120. A representation of the model-predicted Bora secondary structure is shown below in pink, with helices represented as rectangles. The helix suggested to interact with the PLK1 'FW' pocket is bordered in black.

**G:** Raw peak intensities within $^{1}$H-$^{15}$N HSQC spectrum at 10 °C for Bora 1-120 (mirrors the $R_2$ result). A representation of the model-predicted Bora secondary structure is shown below in pink, with helices represented as rectangles. The helix suggested to interact with the PLK1 'FW' pocket is bordered in black.

**H:** Peak intensity ratios to compare the change in intensities with increasing temperature. A representation of the model-predicted Bora secondary structure is shown below in pink, with helices represented as rectangles. The helix suggested to interact with the PLK1 'FW' pocket is bordered in black.

Fig. S9

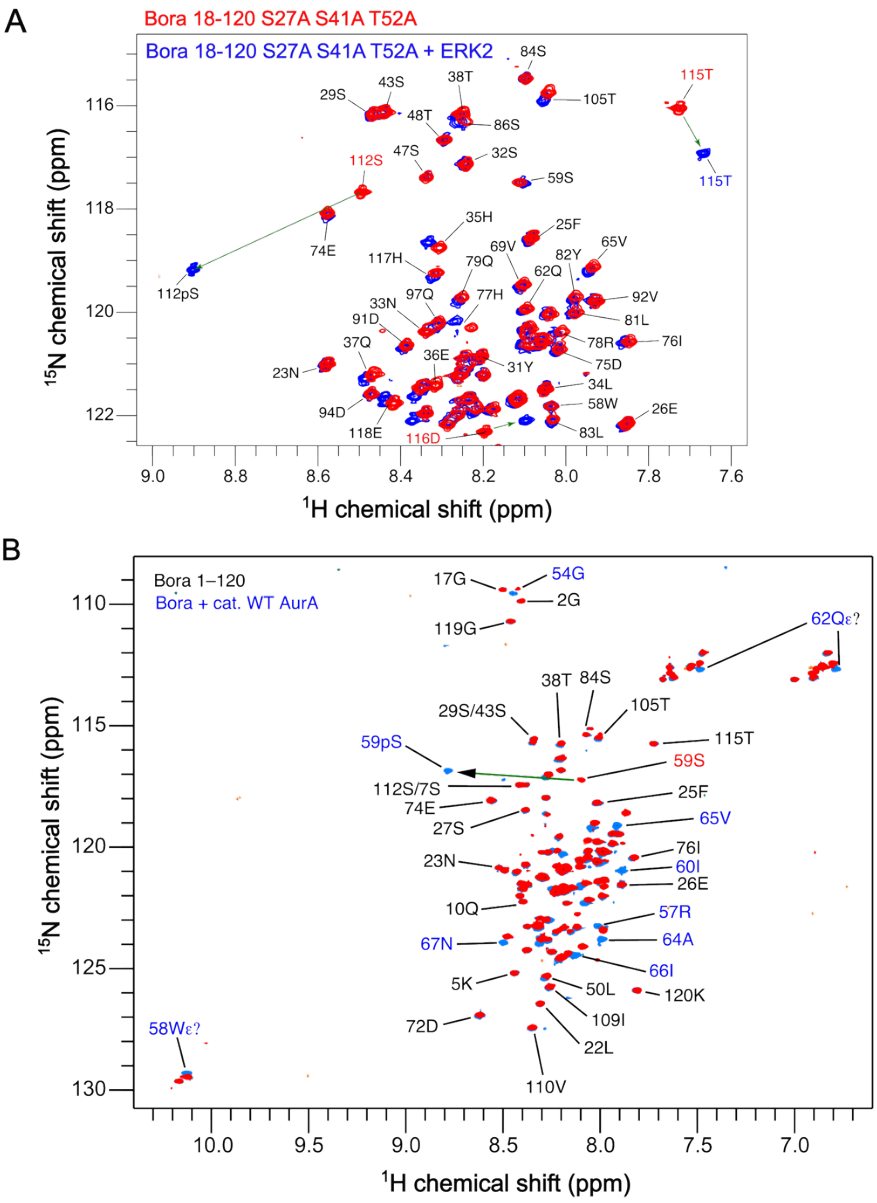


**Figure S9 – Phosphorylation of Bora by Aurora-A and ERK2**

**A:** $^{1}H$-$^{15}N$ HSQC spectra of Bora 18-120 S27A S41A T52A before (red) and after incubation with ERK1 (blue) in the presence of ATP/$Mg^{2+}$. A significant downfield shift is seen for the Ser112 peak, as well as small changes to neighbouring residues, indicating it has been phosphorylated.

**B:** $^{1}H$-$^{15}N$ HSQC spectra of Bora 1-120 before (red) and after incubation with Aurora-A kinase domain (blue) in the presence of ATP/$Mg^{2+}$. A significant downfield shift is seen for the Ser59 peak, as well as small changes to neighbouring residues, indicating it has been phosphorylated.

Fig. S10

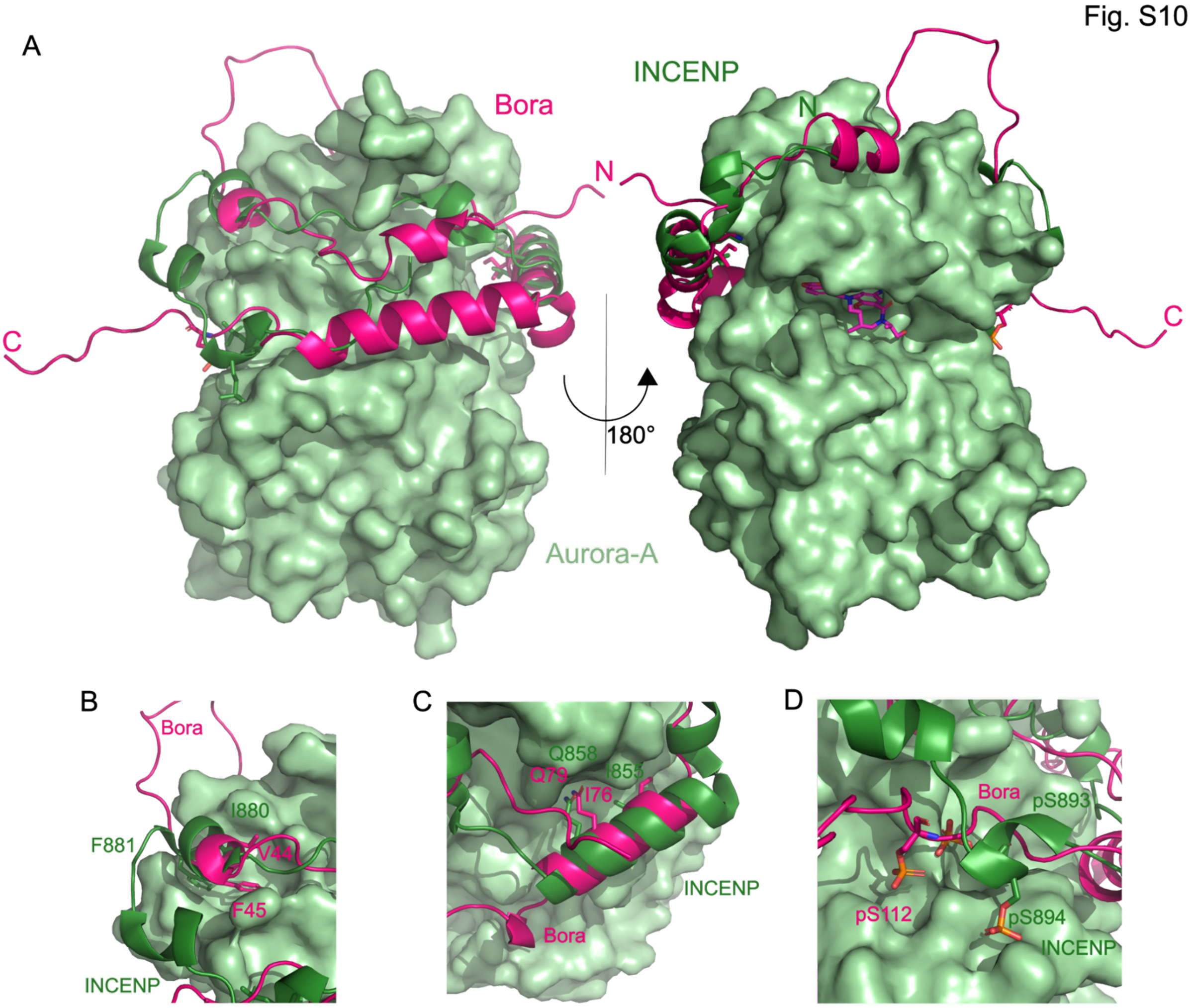


**Figure S10 – Comparison of the predicted Bora:Aurora-A structure with INCENP bound to Aurora-C**

**A:** Comparison of the model of Bora (magenta) bound to Aurora-A (green) with the crystal structure of INCENP bound to Aurora-C (6GR8). INCENP is shown in dark green.

**B:** Comparison of the model of Bora (magenta) bound to Aurora-A (green) with the crystal structure of INCENP bound to Aurora-C (6GR8), focusing on the Y-pocket of Aurora-A. INCENP is shown in dark green. Bora F45 overlaps with where INCENP F881 binds into a comparable pocked on Aurora-C.

**C:** Comparison of the model of Bora (magenta) bound to Aurora-A (green) with the crystal structure of INCENP bound to Aurora-C (6GR8), focusing on the helical region of Bora between 71-85. INCENP is shown in dark green. INCENP is also helical in this region. Both proteins present an isoleucine and glutamine towards the surface of their Aurora binding partner.

**D:** Comparison of the model of Bora (magenta) bound to Aurora-A (green) with the known structure of INCENP bound to Aurora-C (6GR8), focusing on the region near the activation loop of Aurora. INCENP is shown in dark green. Phosphorylated Ser112 is modelled near to where phosphorylated INCENP binds to Aurora-C.

Fig S11.

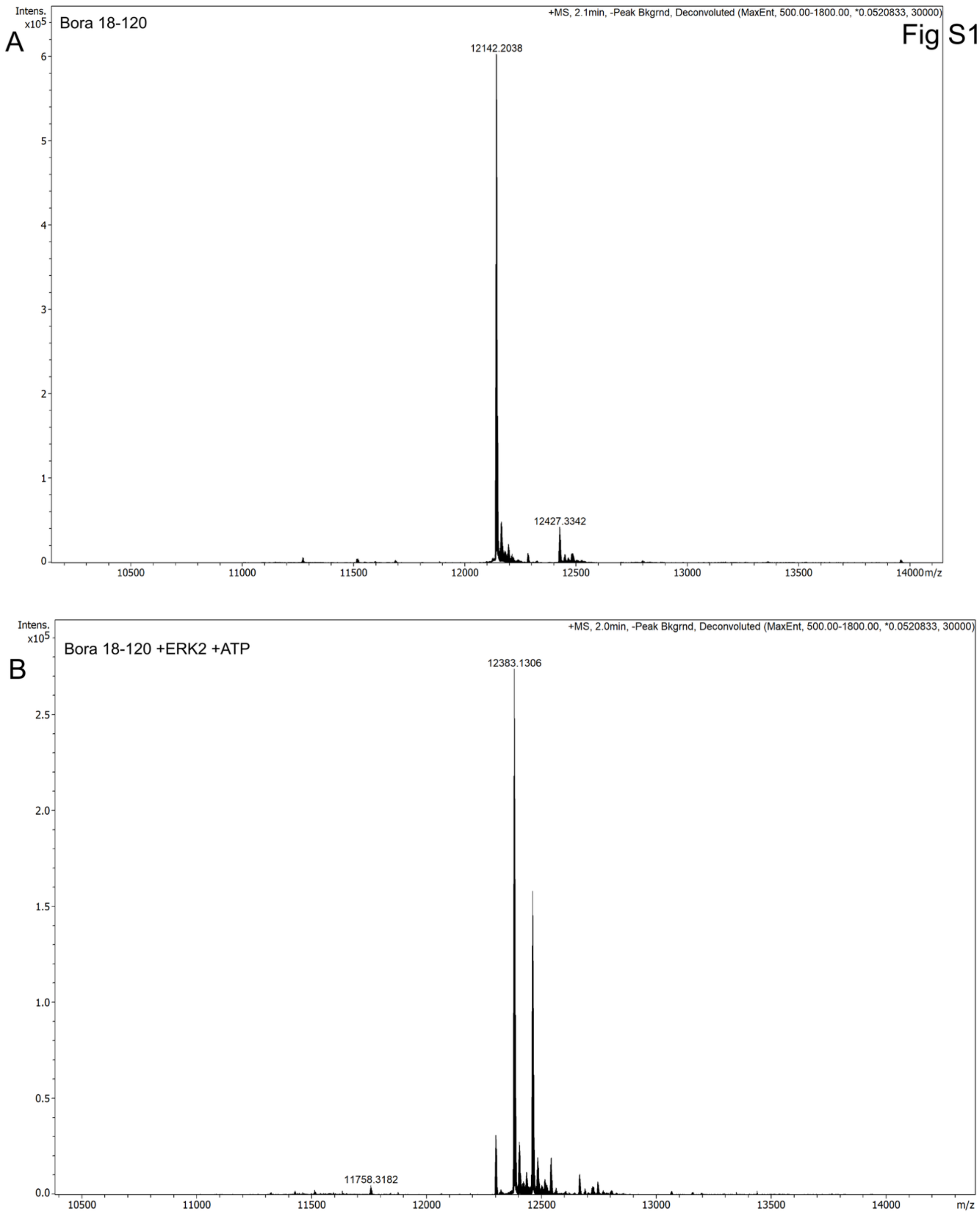
A
Bora 18-120
+MS, 2.1min, -Peak Bkgrnd, Deconvoluted (MaxEnt, 500.00-1800.00, *0.0520833, 30000)
Intens. x10^5
12142.2038
12427.3342
m/z
B
Bora 18-120 +ERK2 +ATP
+MS, 2.0min, -Peak Bkgrnd, Deconvoluted (MaxEnt, 500.00-1800.00, *0.0520833, 30000)
Intens. x10^5
12383.1306
11758.3182
m/z

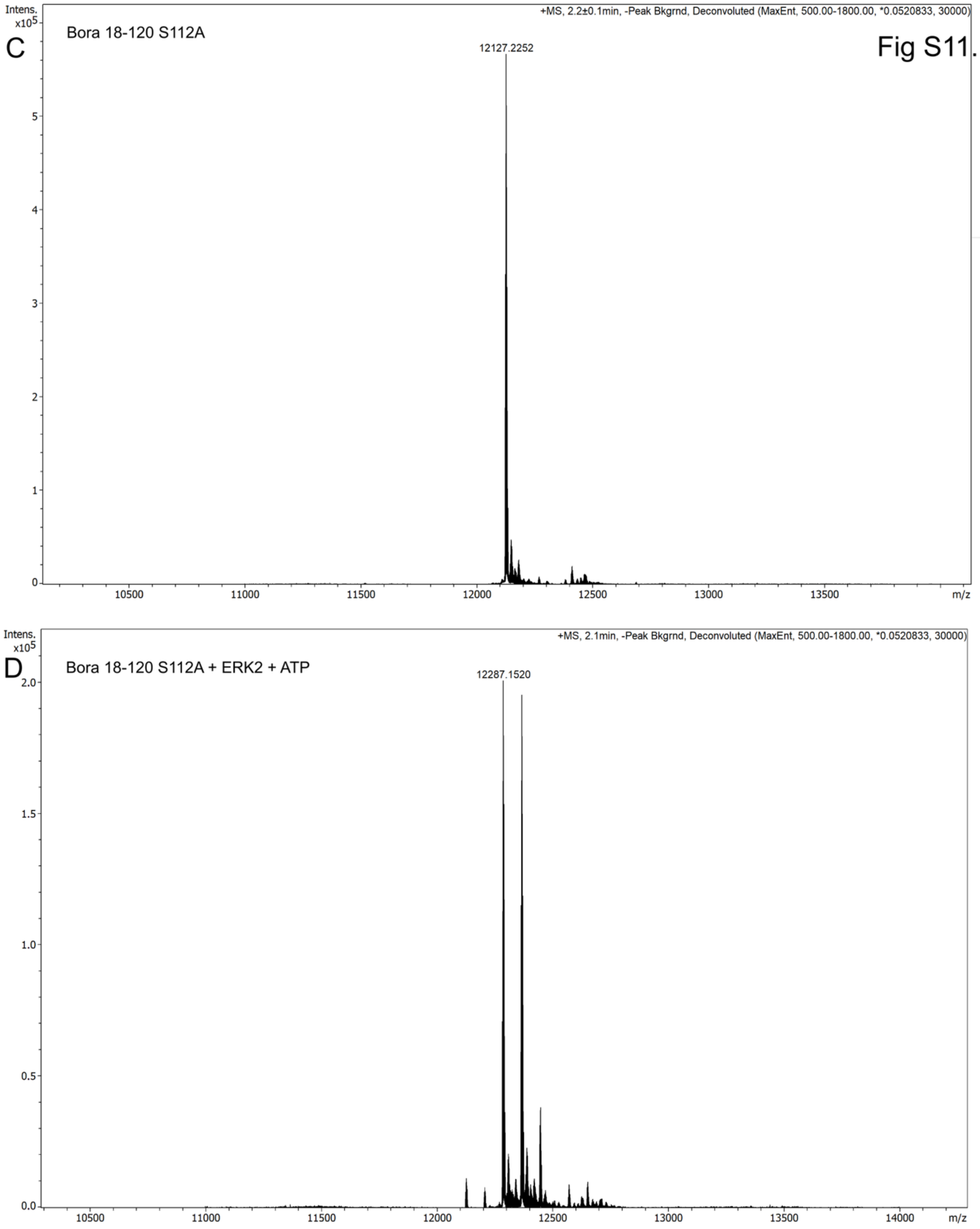
Fig S11.
C
Bora 18-120 S112A
+MS, 2.2±0.1min, -Peak Bkgrnd, Deconvoluted (MaxEnt, 500.00-1800.00, *0.0520833, 30000)
Intens. x10^5
12127.2252
m/z
D
Bora 18-120 S112A + ERK2 + ATP
+MS, 2.1min, -Peak Bkgrnd, Deconvoluted (MaxEnt, 500.00-1800.00, *0.0520833, 30000)
Intens. x10^5
12287.1520
m/z

Fig S11.

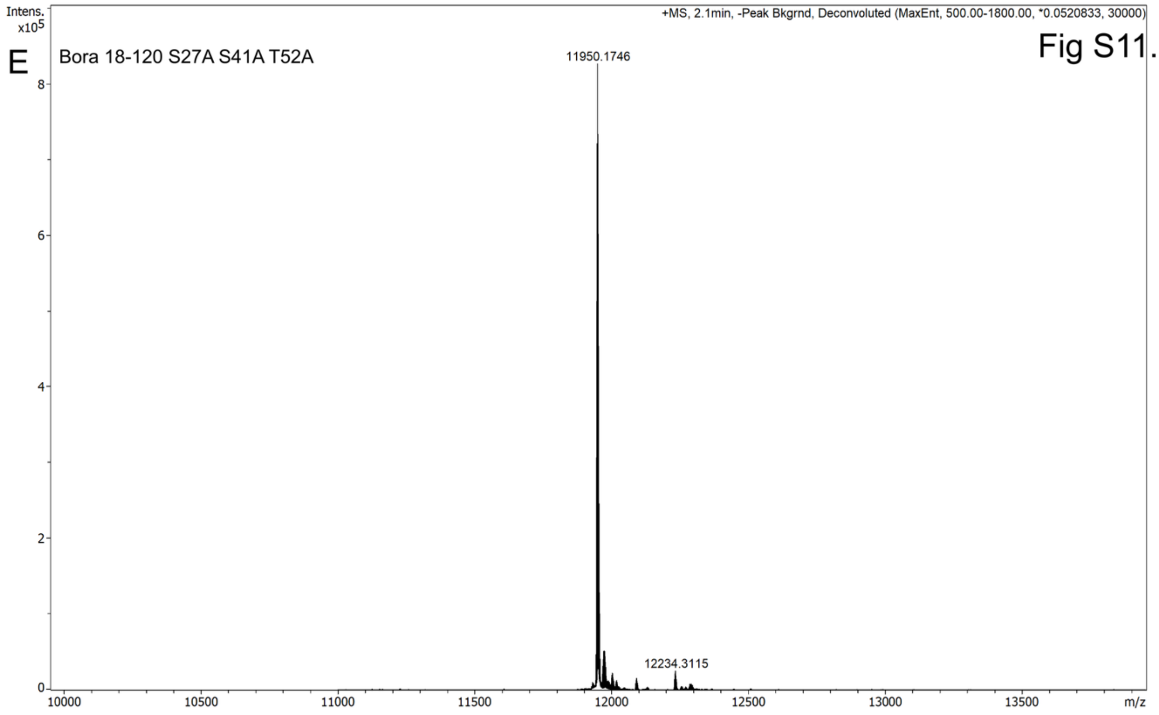

E
Bora 18-120 S27A S41A T52A
+MS, 2.1min, -Peak Bkgrnd, Deconvoluted (MaxEnt, 500.00-1800.00, *0.0520833, 30000)
Intens. x10^5
11950.1746
12234.3115
m/z


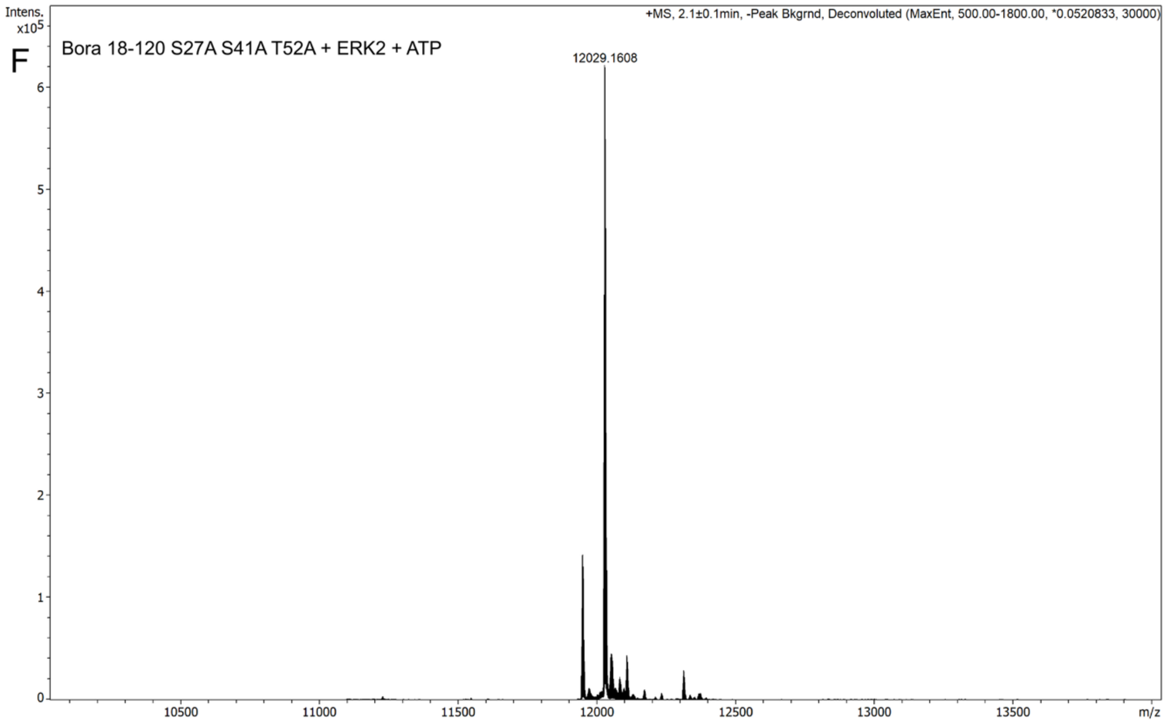

F
Bora 18-120 S27A S41A T52A + ERK2 + ATP
+MS, 2.1±0.1min, -Peak Bkgrnd, Deconvoluted (MaxEnt, 500.00-1800.00, *0.0520833, 30000)
Intens. x10^5
12029.1608
m/z

Fig S11.

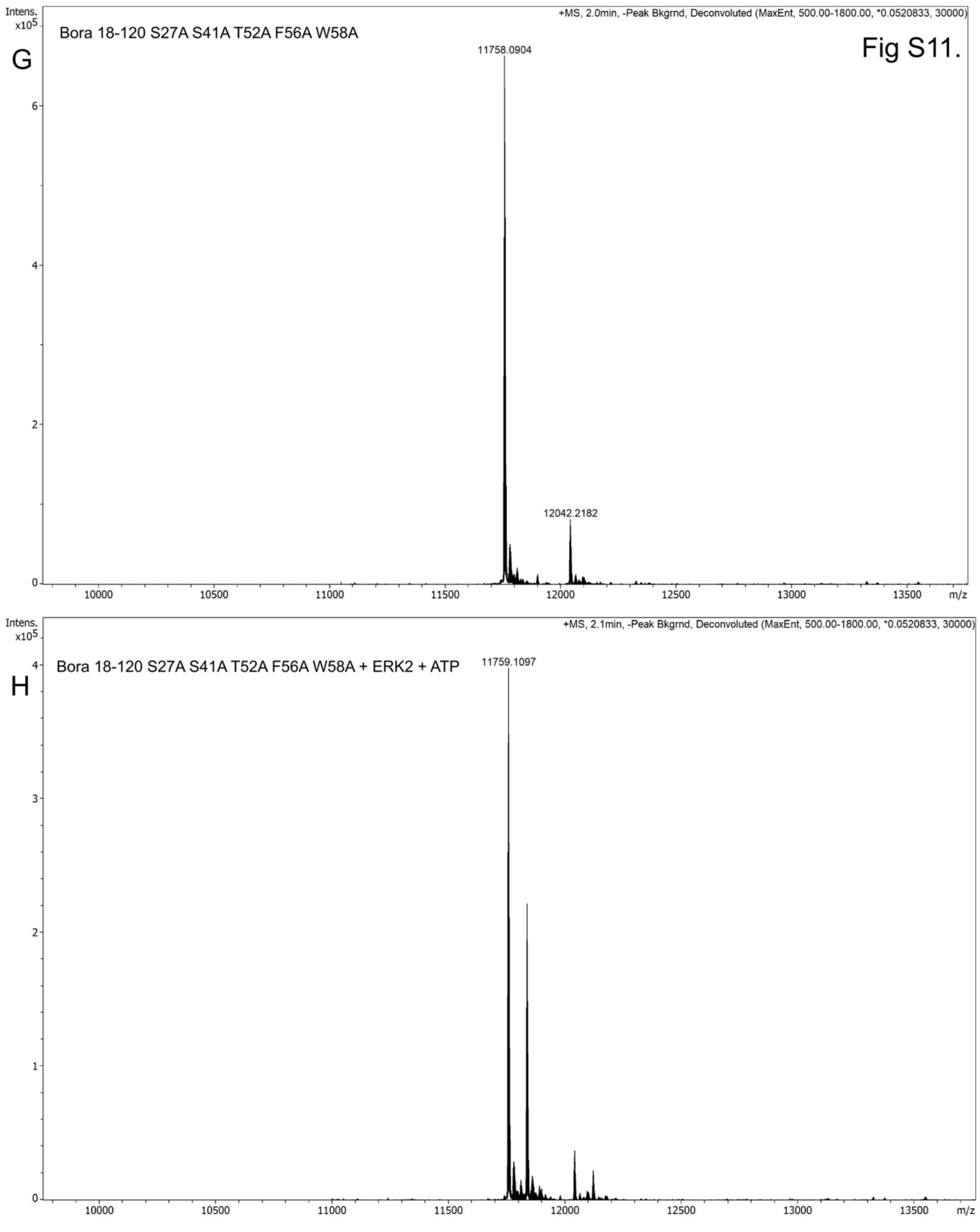
G
Bora 18-120 S27A S41A T52A F56A W58A
+MS, 2.0min, -Peak Bkgrnd, Deconvoluted (MaxEnt, 500.00-1800.00, *0.0520833, 30000)
Intens. x10^5
11758.0904
12042.2182
m/z
H
Bora 18-120 S27A S41A T52A F56A W58A + ERK2 + ATP
+MS, 2.1min, -Peak Bkgrnd, Deconvoluted (MaxEnt, 500.00-1800.00, *0.0520833, 30000)
Intens. x10^5
11759.1097
m/z

Fig S11.

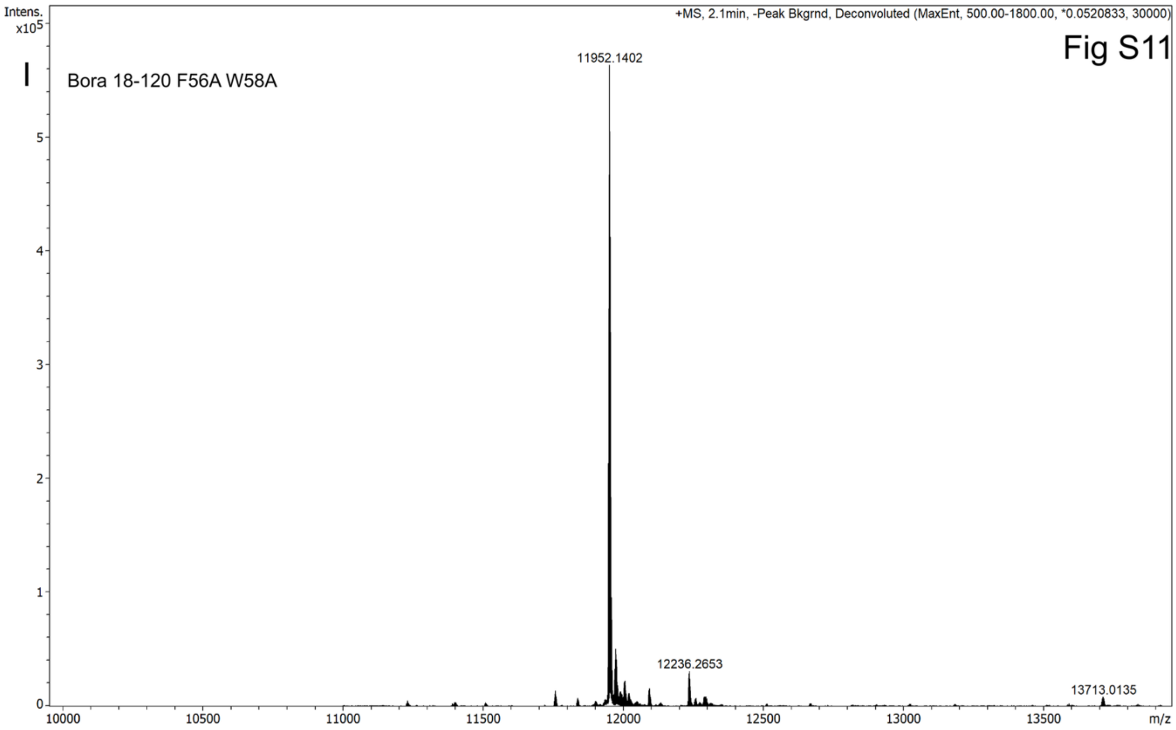
Intens.
x10^5
+MS, 2.1min, -Peak Bkgrnd, Deconvoluted (MaxEnt, 500.00-1800.00, *0.0520833, 30000)
I
Bora 18-120 F56A W58A
11952.1402
12236.2653
13713.0135
m/z

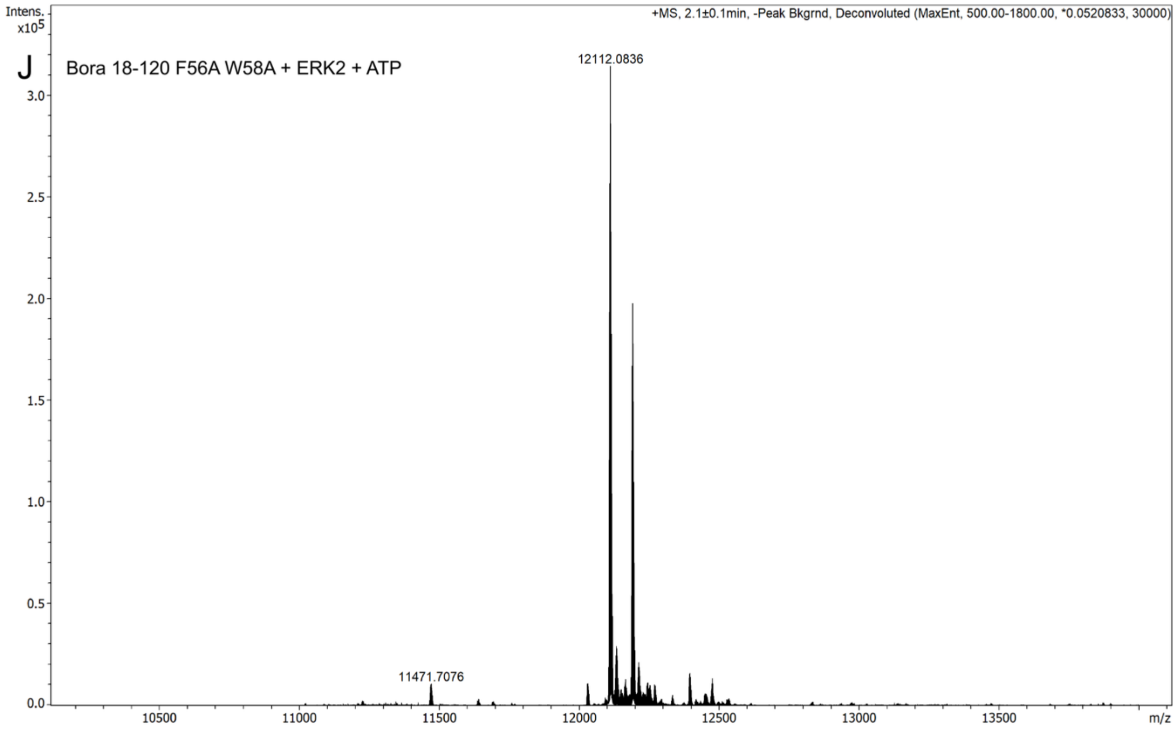
Intens.
x10^5
+MS, 2.1±0.1min, -Peak Bkgrnd, Deconvoluted (MaxEnt, 500.00-1800.00, *0.0520833, 30000)
J
Bora 18-120 F56A W58A + ERK2 + ATP
12112.0836
11471.7076
m/z

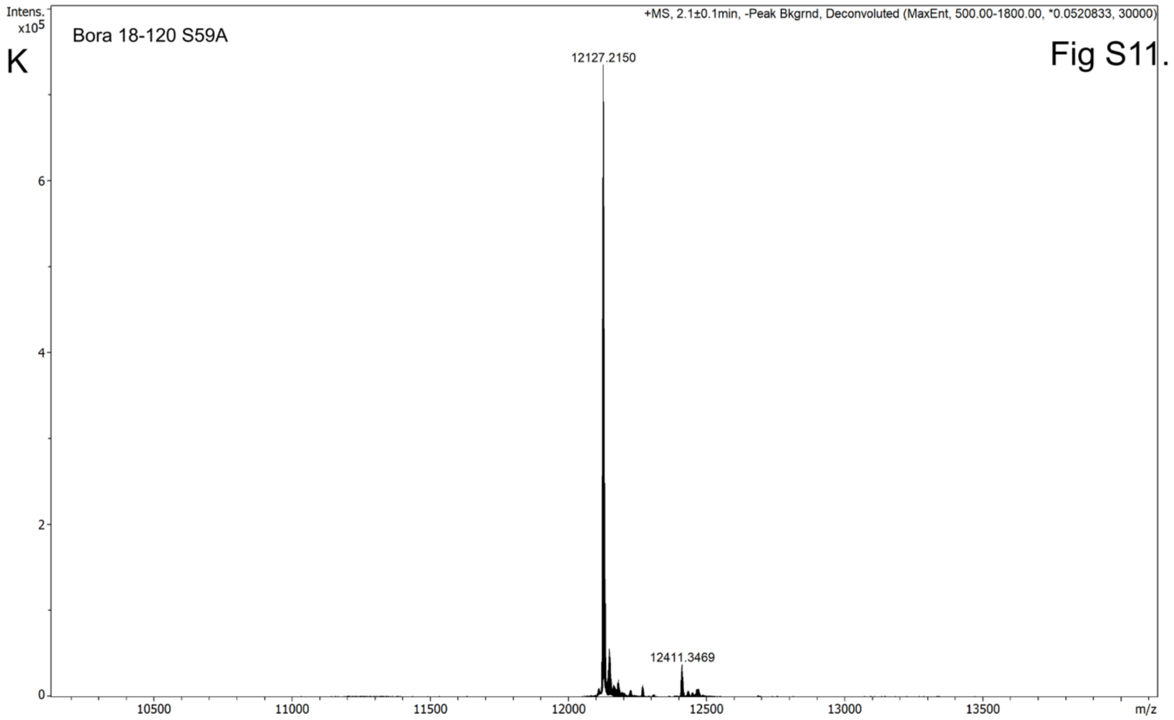
Intens.
x10^5
K
Bora 18-120 S59A
+MS, 2.1±0.1min, -Peak Bkgrnd, Deconvoluted (MaxEnt, 500.00-1800.00, *0.0520833, 30000)
Fig S11.
12127.2150
12411.3469
m/z

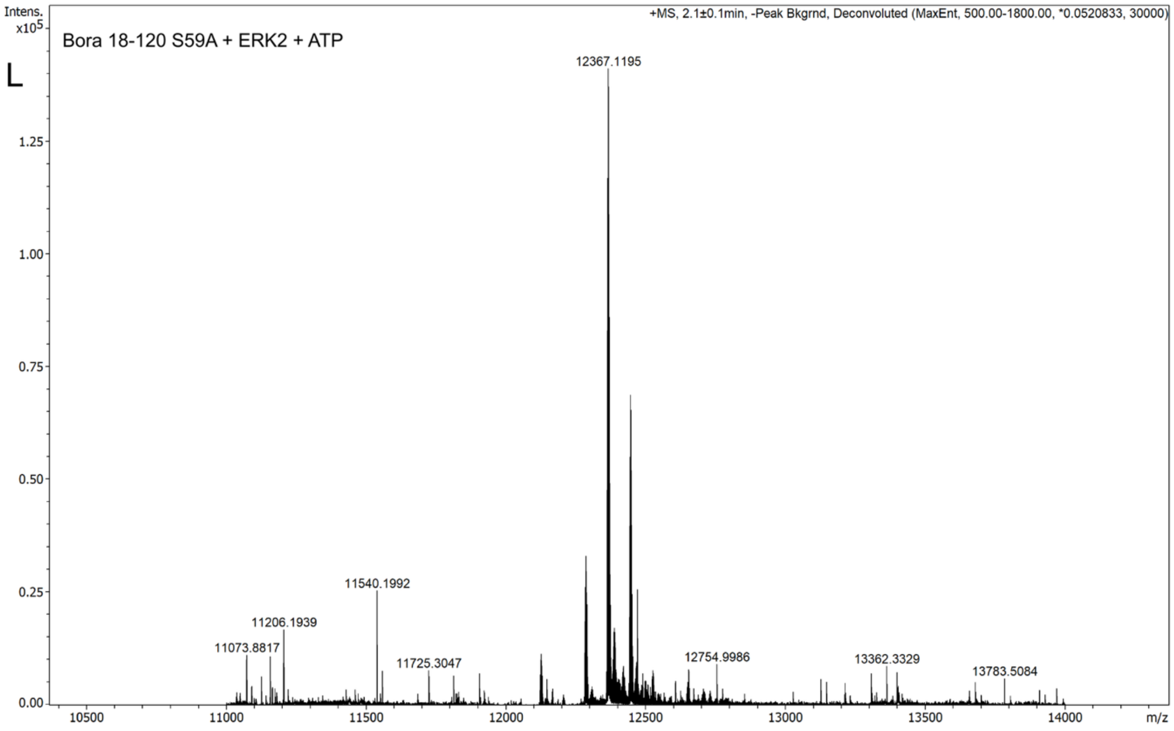
Intens.
x10^5
L
Bora 18-120 S59A + ERK2 + ATP
+MS, 2.1±0.1min, -Peak Bkgrnd, Deconvoluted (MaxEnt, 500.00-1800.00, *0.0520833, 30000)
12367.1195
11540.1992
11206.1939
11073.8817
11725.3047
12754.9986
13362.3329
13783.5084
m/z

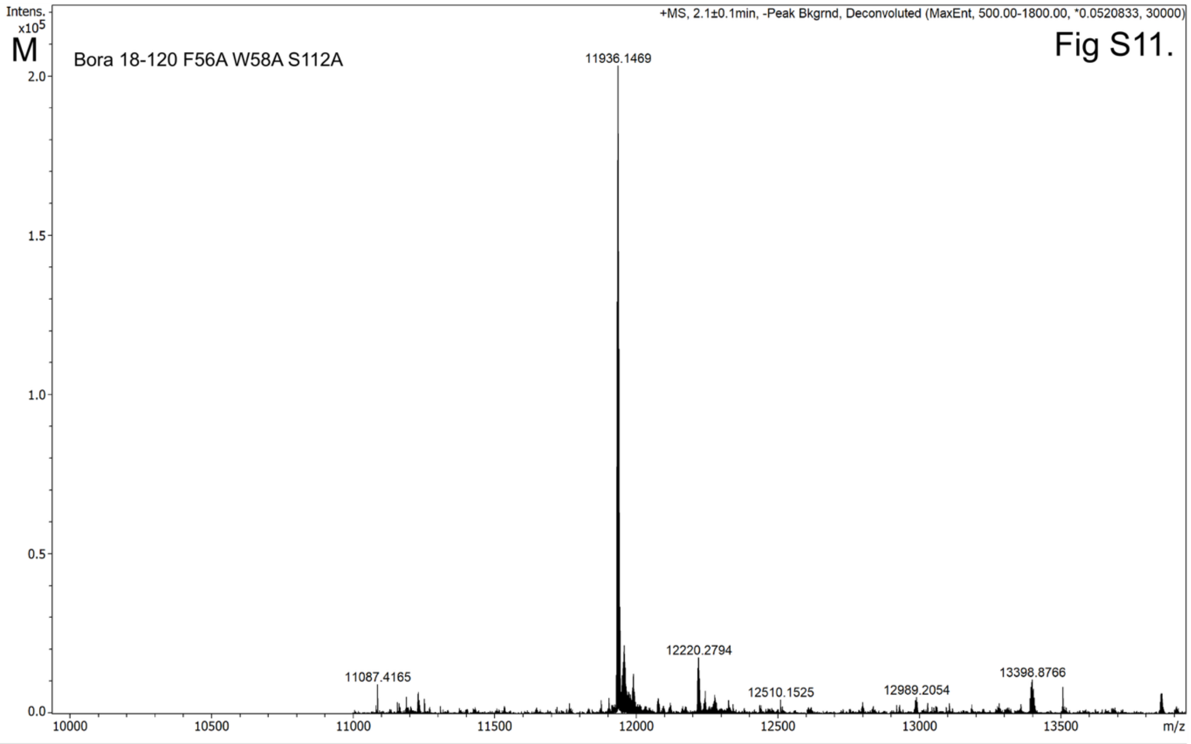


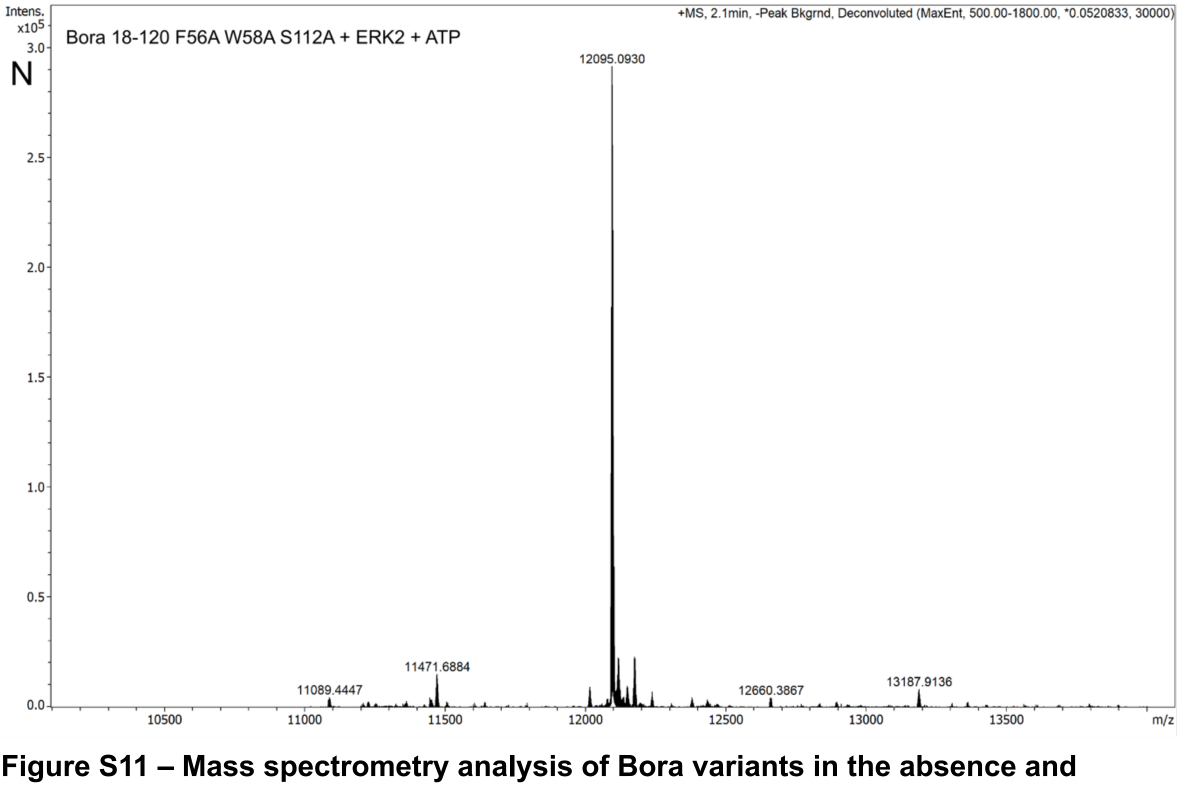


**Figure S11 – Mass spectrometry analysis of Bora variants in the absence and presence of ERK2.**

**A:** Mass spectrometry analysis of wild-type Bora 18-120.

**B:** Mass spectrometry analysis of wild-type Bora 18-120 after a 1 hour incubation with ERK2 and ATP before quenching with EDTA.

**C:** Mass spectrometry analysis of Bora S112A 18-120.

**D:** Mass spectrometry analysis of Bora S112A 18-120 after a 1 h incubation with ERK2 and ATP before quenching with EDTA.

**E:** Mass spectrometry analysis of Bora S27A S41A T52A 18-120.

**F:** Mass spectrometry analysis of Bora S27A S41A T52A 18-120 after a 1 h incubation with ERK2 and ATP before quenching with EDTA.

**G:** Mass spectrometry analysis of Bora S27A S41A T52A F56AW58A 18-120.

**H:** Mass spectrometry analysis of Bora S27A S41A T52A F56AW58A 18-120 after a 1 h incubation with ERK2 and ATP before quenching with EDTA.

**I:** Mass spectrometry analysis of Bora F56AW58A 18-120.

**J:** Mass spectrometry analysis of Bora F56AW58A 18-120 after a 1 h incubation with ERK2 and ATP before quenching with EDTA.

**K:** Mass spectrometry analysis of Bora S59A 18-120.

**L:** Mass spectrometry analysis of Bora 18-120 S59A after a 1 h incubation with ERK2 and ATP before quenching with EDTA.

**M:** Mass spectrometry analysis of Bora F56AW58A S112A 18-120.

**N:** Mass spectrometry analysis of Bora F56AW58A S112A 18-120 after a 1 h incubation with ERK2 and ATP before quenching with EDTA .

**Table S1**. Primers used in site directed mutagenesis

| Construct | Mutation | Forward primer | Reverse primer |
|---|---|---|---|
| Human PLK1 3-330 | K82R | GTTCGCGGGCaggATTGTGCCTA | ACCTCCTTGGTGTCCGCG |
| | R106A | ATCCATTCACgccAGCCTCGCCC | ATTTCCATAGACATCTTCTCC |
| | S99R | GGAGAAGATGcgtATGGAAATATCCATTCACCGCAG | CTCTGGTGCGGCTTGAGC |
| Human Bora 18-120 | S112A | GATCGTTCCGGCGCCGTGGACCG | ACGTCTTTGGTAAAGAATTC |
| | F56AW58A | tgctAGCATTGATCAGCTGGCGGTG | cgagcCTTGCCCGGGGTCGGCAG |
| | S59A | GTTTCGTTGGgcgATTGACCAGCTGGCGGTGATCAAC | TTGCCCGGGGTCGGC |

**Table S2**. Constructs used within this study

| | Vector | Mutations |
|---|---|---|
| PLK1 3-330 | pETSUMO | K82R |
| | | K82R S99R R106A |
| Aurora-A 122-403 | pET30TEV | |
| | pET30TEV | C290A C393A |
| Bora 1-120 | pETSUMO | |
| Bora 18-120 | pETSUMO | |
| | pETSUMO | S59A |
| | pETSUMO | S112A |
| | pETSUMO | F56AW58A |
| | pETSUMO | F56A W58A S112A |
| | pET28a+ | S27A S41A T52A F56A W58A |
| | pET28a+ | S27A S41A T52A |
| CEP192 442-533 | pETSUMO | |
| TPX2 1-43 | pGEX-cs | |
| DARPin | pET28a+ | |

**Table S3**. iPTM scores from AlphaFold2 and AlphaFold3 predictions

| **Aurora-A** | **Bora** | **PLK1** | **Top ipTM AlphaFold2** | **Top ipTM AlphaFold3** |
|---|---|---|---|---|
| 1-403 | 1-559 | 1-603 | 0.63 | 0.65 |
| 1-403 | 1-559 | | 0.67 | 0.64 |
| 122-403 | 18-120 | | 0.82 | 0.83 |
| 122-403 | 18-120 | 30-330 | 0.77 | 0.77 |
| | 1-559 | 1-603 | 0.57 | 0.61 |
| | 18-120 | 1-603 | 0.36 | 0.51 |
| 1-403 | | 1-603 | 0.27 | 0.21 |

**Table S4**. Mass spectrometry analysis of modified Bora samples

| **Fig** | **Bora version** | **Major species** | **Other species** |
|---|---|---|---|
| S11A | WT 18-120 (**Calc. mw 12143**) | 12142 (100%) | |
| S11B | WT 18-120 + ERK2 + ATP | 12383 (65%) (3P) | 12463 (30%) (4P)<br>12303 (5%) (2P) |
| S11C | S112A 18-120 (**Calc. mw 12127**) | 12127 (100%) | |
| S11D | S112A 18-120 + ERK2 + ATP | 12287 (45%) (2P) | 12367 (45%) (3P)<br>12447 (10%) (4P) |
| S11E | S27A S41A T52A 18-120 (**Calc. mw 11950**) | 11950 (100%) | |
| S11F | S27A S41A T52A 18-120 + ERK2 + ATP | 12029 (85%) (1P) | 11950 (15%) (0P) |
| S11G | S27A S41A T52A F56AW58A 18-120 (**Calc. mw 11759**) | 11758 (100%) | |
| S11H | S27A S41A T52A F56AW58A 18-120 + ERK2 + ATP | 11758 (65%) (0P) | 11838 (35%) (1P) |
| S11I | F56AW58A 18-120 (**Calc. mw 11952**) | 11952 (100%) | |
| S11J | F56AW58A 18-120 + ERK2 + ATP | 12112 (65%) (2P) | 12192 (35%) (3P) |
| S11K | S59A 18-120 (**Calc. mw 12127**) | 12127 (100%) | |
| S11L | S59A 18-120 + ERK2 + ATP | 12367 (60%) (3P) | 12447 (30%) (4P)<br>12287 (10%) (2P) |
| S11M | F56AW58A S112A 18-120 (**Calc. mw 11936**) | 11936 (100%) | |
| S11N | F56AW58A S112A 18-120 + ERK2 + ATP | 12095 (100%) (2P) | |
